# Ultrastructural diversity and subcellular organization of nigral Lewy pathology in Parkinson’s disease

**DOI:** 10.1101/2024.07.25.605088

**Authors:** Amanda J Lewis, Lukas van den Heuvel, Marta di Fabrizio, Kaliya Bandelier, Daria Proniakova, Domenic Burger, Notash Shafiei, Kazadi Ekundayo, Selene Offringa, Evelien Huisman, John GJM Bol, Wilma DJ Van de Berg, Henning Stahlberg

## Abstract

Lewy bodies, the defining pathological feature of Parkinson’s disease, are intraneuronal inclusions enriched in aggregated alpha-synuclein (αSyn). We used correlative light and electron microscopy to selectively investigate phosphorylated αSyn (αSyn^pS129^)-positive inclusions in the *substantia nigra* of end-stage postmortem Parkinson’s disease brain. Here we show that somatic αSyn^pS129^ inclusions in nigral dopaminergic neurons are consistently fibrillar, whereas the membranous-type inclusions are restricted to neuritic processes. These neuritic inclusions displayed marked ultrastructural heterogeneity, ranging from predominantly membranous to mixed membranous-fibrillar forms. The selective targeting of defined inclusions enabled detailed structural characterization of Lewy pathology, rather than quantitative or disease-stage comparisons. Our findings highlight clear ultrastructural differences between somatic and neuritic αSyn^pS129^ pathology and demonstrate the structural complexity and heterogeneity of Lewy pathology in human Parkinson’s disease brain.

## Introduction

Lewy bodies (LBs) are abnormal intracellular inclusions rich in aggregated alpha-synuclein (αSyn). First identified in the *substantia nigra*, where defects in the dopaminergic neurons lead to the characteristic motor function failures in PD patients^1^, they now represent the major pathological hallmark of Parkinson’s disease (PD) and dementia with Lewy bodies. Classical histopathology of post-mortem brain tissue described LBs as spherical structures with a dense core and a pale halo^2^ that was immunopositive for αSyn^3^. Subsequent transmission electron microscopy (EM) attributed the core to highly electron-dense material, and the halo to radially arranged αSyn-fibrils^4–7^.

Other forms of Lewy pathology, including Lewy neurites/threads, bulgy neurites and pale bodies, have also been reported to contain αSyn fibrils as a major component. Lewy threads are thin, elongated deposits of αSyn along axons and dendrites, bulgy neurites are characterized by localized rod-shaped swellings along neuritic processes^8^, and pale bodies are large cytoplasmic aggregates, that are weakly stained by hematoxylin and eosin^9^, often irregular in shape and with an ultrastructure consisting of randomly organized fibrils intermixed with clusters of mitochondria and other organelles^9–11^.

The predominance of fibrillar ultrastructures, together with experimental evidence from model systems, has led to the prion-like model of Lewy pathology formation, in which αSyn misfolds, assembles into oligomers and fibrils, and progressively accumulates into inclusions^12–15^. However, increasing evidence indicates that fibril formation represents only one component of a broader spectrum of αSyn aggregation states. Recent high-end microscopy and -omics studies showed that LBs contain lipids^16,17^, organelles^5^, membranous fragments^18^, and more than 100 additional proteins^7,19^. Additionally, our recent correlative light and electron microscopy (CLEM) study characterized other forms of αSyn pathology enriched in lipids and membranous structures^18^. These structures, here referred to as membranous pathology, are ultrastructurally distinct from fibrillar inclusions by their high proportion of vesicles and membrane fragments co-localizing with αSyn immunoreactivity in the absence of any observable fibrils. At the molecular scale, cryo-electron microscopy of αSyn fibrils extracted from human PD and DLB brain has revealed distinct conformational strains^20,20,21^, highlighting structural polymorphism within fibrillar assemblies.

The molecular diversity of αSyn pathology is paralleled by heterogeneity in antibody labeling, with numerous microscopy studies reporting differential αSyn epitope staining patterns across synucleinopathies^22–29^. Multiple post-translational modifications including phosphorylation, ubiquitination and truncation of the C-terminus are believed to modulate αSyn misfolding and LB formation(reviewed in^30^). In particular, the finding that αSyn in LBs is extensively phosphorylated at the serine residue 129 (αSyn^pS129^)^31^ paved the way for its use as a key biochemical and histological marker for synucleinopathies.

Several human post-mortem studies have proposed a morphological sequence for LB formation, based on the different morphologies of αSyn immuno-staining in combination with disease staging^8,32–34,22^. This proposed process includes the accumulation of diffuse cytoplasmic staining or punctate dots, followed by the development of a small aggregation center that grows into a pale body and finally develops into the halo LB. Pale bodies are considered precursors to LBs, are more abundant in the early stages of PD compared to halo LBs, and several pale bodies often co-exist in the same cell^9,35^. The appearance of Lewy neurites is thought to precede the appearance of somal αSyn pathology^8^, yet it is unclear how Lewy neurites specifically relate to the development of the LBs. Further, it is not known how the membranous pathology fits into the proposed timeline for LB formation. As there are currently no animal or cellular models that can reproduce the large diversity of human PD pathology, to infer a theory on the mechanism for LB formation requires studying their morphological heterogeneity in the human post-mortem brain.

In this study, we used CLEM to investigate αSyn^pS129^ inclusions in the *substantia nigra* of post-mortem PD brain donors, targeting a large diversity of morphologies. By directly correlating immunofluorescence with ultrastructure, we aimed to comprehensively resolve the subcellular organization and morphological diversity of Lewy pathology in human brain. By imaging 144 αSyn^pS129^ inclusions from the *substantia nigra* of 12 brain donors we found that the pathology can be classified based on distinct ultrastructural characteristics, depending upon whether it occurs in the soma of dopaminergic neurons or not. Somal inclusions consistently exhibited fibrillar ultrastructures. The increasing density of accumulated fibrils matches previously proposed timelines for LB formation^22,32^ culminating in three distinct stages of fibril organization for halo LBs. Notably, all membranous inclusions were observed in neuritic compartments and displayed a variety of ultrastructures including membranous only (with an absence of fibrils), membranes intermixed with fibrils, and a halo of membranes surrounding a fibrillar core. We present an example where all these ultrastructures are present within the same neuritic inclusion suggesting that membranous inclusions may provide the environment for the formation of αSyn fibrils. Our analysis reveals a clear subcellular segregation between fibrillar and membranous αSyn^pS129^ inclusions and provides an ultrastructural framework that refines current models of Lewy body formation. These findings extend previous observations of membrane-associated pathology and offer insight into how different forms of αSyn aggregation relate to neuronal compartmentalization in PD.

## Results

### Localization of αSyn pathology using CLEM

To localize pathology in post-mortem human brain, we performed CLEM on free-floating 40-60 μm semi-thin sections immuno-labeled against αSyn^pS129^. Pathology was identified either by fluorescence microscopy prior to embedding (CLEM^FL^) or post-embedding using immunohistochemistry (IHC; CLEM^IHC^)^36^. In free-floating sections, we targeted pathology in dopaminergic neurons in the *substantia nigra* (defined by presence of neuromelanin granules) showing various staining patterns by immuno-fluorescence. These included diffuse/punctate cellular αSyn^pS129^, small (< 5μm) and large (> 5 μm) inclusions with diffuse αSyn^pS129^ staining, inclusions with a distinct αSyn^pS129^ halo (halo LBs), and diffuse αSyn^pS129^ inclusions with halo LBs at the periphery. A neurofilament marker (against neurofilament heavy chain) and DAPI staining for cell nuclei were used to help delineate the cellular context for the inclusions where neuromelanin was absent (Figure 1a-e).

**Figure 1.**
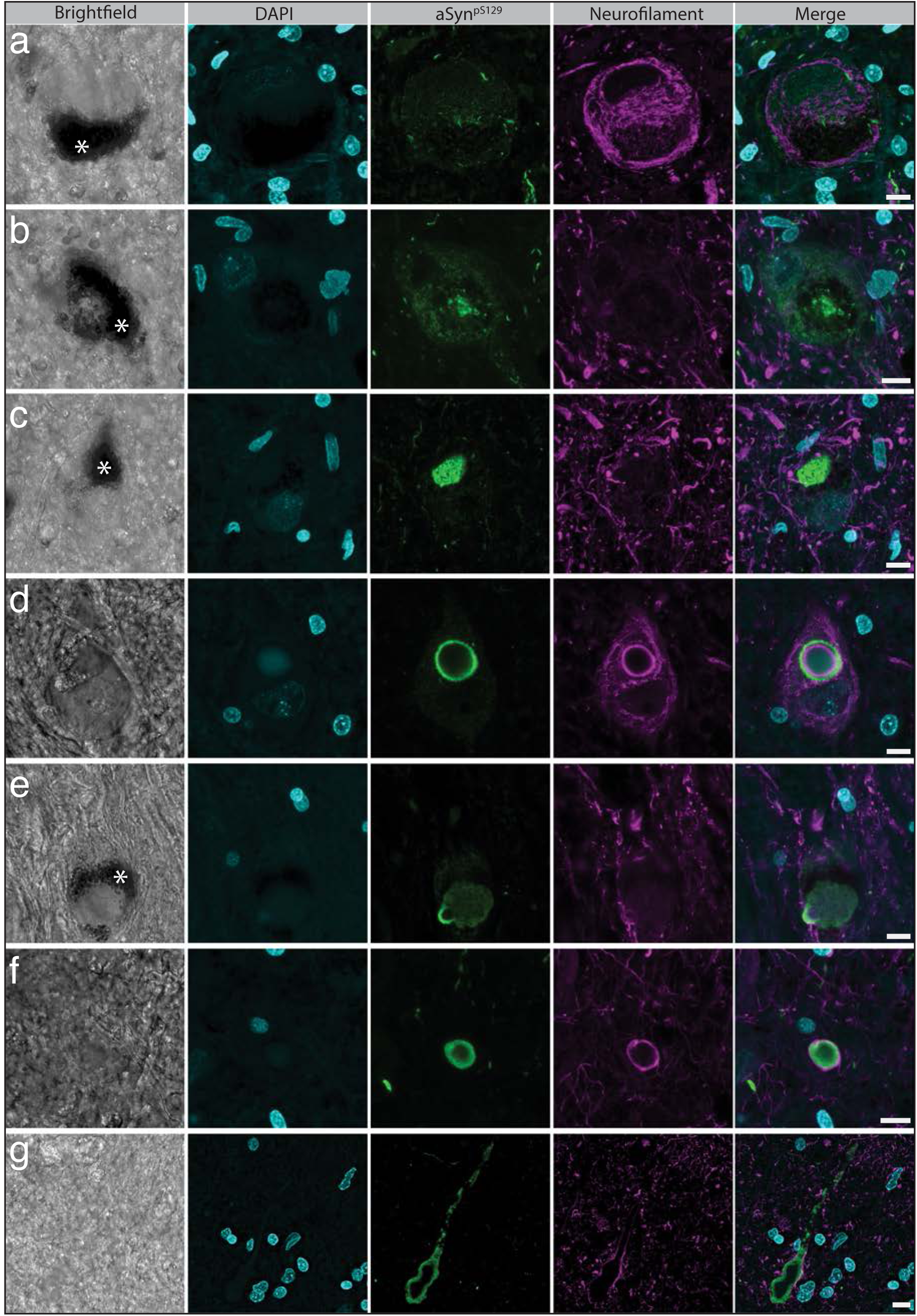
Morphological staining patterns of αSyn^pS129^ pathology targeted for EM. Maximum projection images of confocal z-stacks of **(a)** diffuse/punctate cellular staining (z = 13.8 μm), **(b)** small aggregates (z = 9 μm) **(c)** large aggregates (z = 9 μm) **(d)** halo LBs (z = 5.4 μm) **(e)** large aggregates with a halo LB at the periphery (z = 7.5 μm) **(f)** compact neuritic aggregates (z = 4.2 μm) and **(g)** Lewy/bulgy neurites (z = 6 μm). The panels in each row show the same field of view. For each panel the brightfield, nuclear staining (DAPI; cyan), αSyn^pS129^(11a5, Prothena; green), neurofilament (neurofilament H, Abcam; magenta) and a merge of the fluorescent channels are shown. The presence of neuromelanin, visible in the brightfield channel (white asterisk), and/or neurofilament and DAPI staining was used to identify somal inclusions (a-e). Scale bars: 10 μm.

Large αSyn^pS129^ aggregates (> 5 μm) that were not clearly within a neuronal cell soma (due to absence of neuromelanin and cellular markers; Figure 1f) as well as bulgy Lewy neurites (Figure 1g) were also included for investigation with CLEM. We often observed a ring of neurofilament staining on the inside of intracellular αSyn^pS129^ halo LBs, consistent with previous reports by Moors et al.^22^ (Figure 1d). A similar neurofilament signal was observed surrounding the non-somal αSyn^pS129^ inclusions (Figure 1f), indicating that these inclusions were located in neuritic compartments (e.g. axon, dendrite or a swollen synapse). As these inclusions were not associated with somatic features and had similar dimensions in the x,y and z planes, their profile suggests a compact neuritic morphology, rather than cross-sections through elongated bulgy Lewy neurites (Figure 1g).

Using this CLEM pipeline, we characterized 146 αSyn^pS129^ immunopositive inclusions from within the *substantia nigra* of twelve brain donors. All inclusions were classified based on their subcellular location and ultrastructural features, and the full morphological classification across donors is summarized in Table 1. Donor information is provided in Supplementary Table 1. In total, 42 representative inclusions are described in detail here, with the full dataset available for download on BioImage Archive.

**Table 1.**
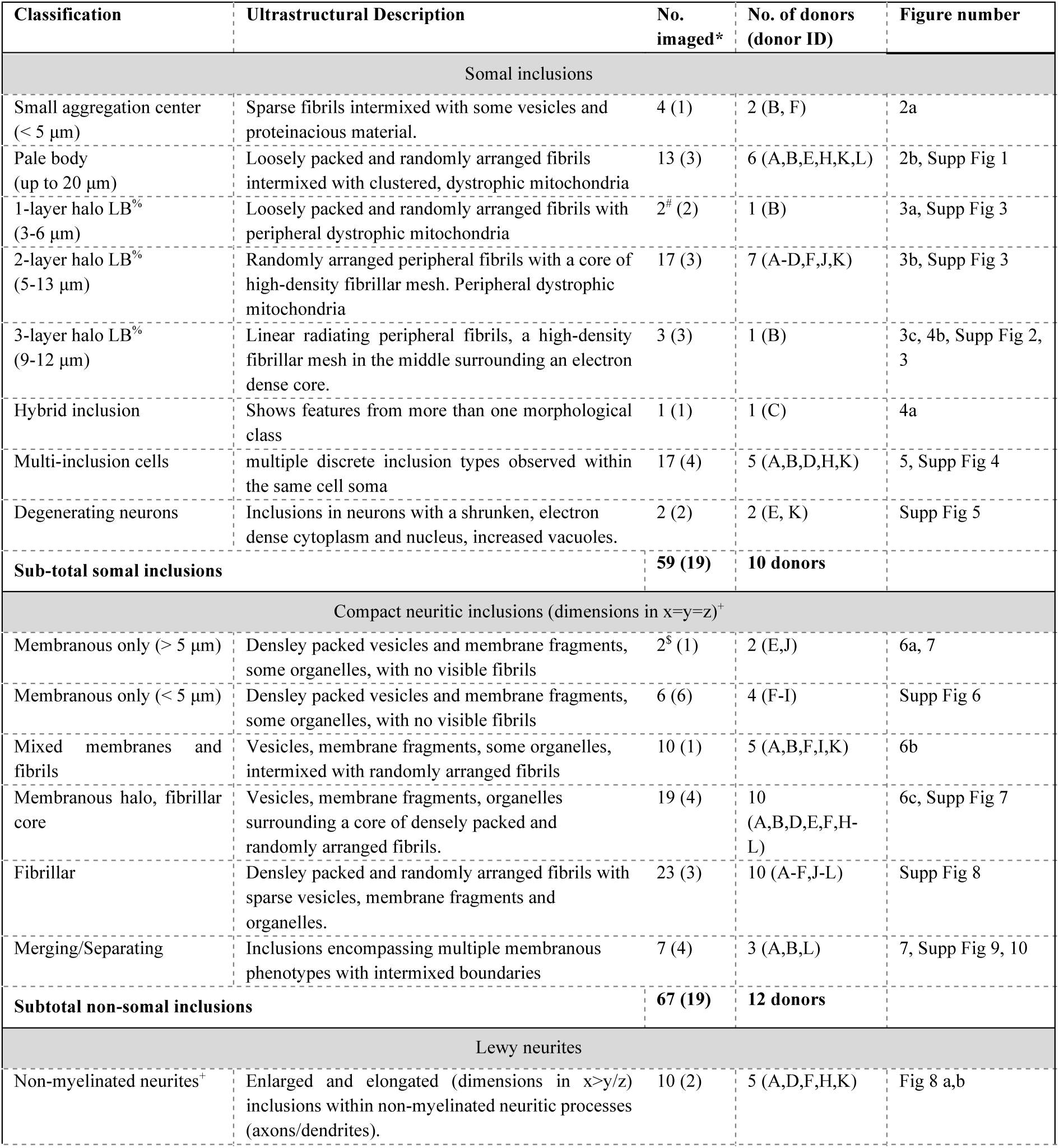
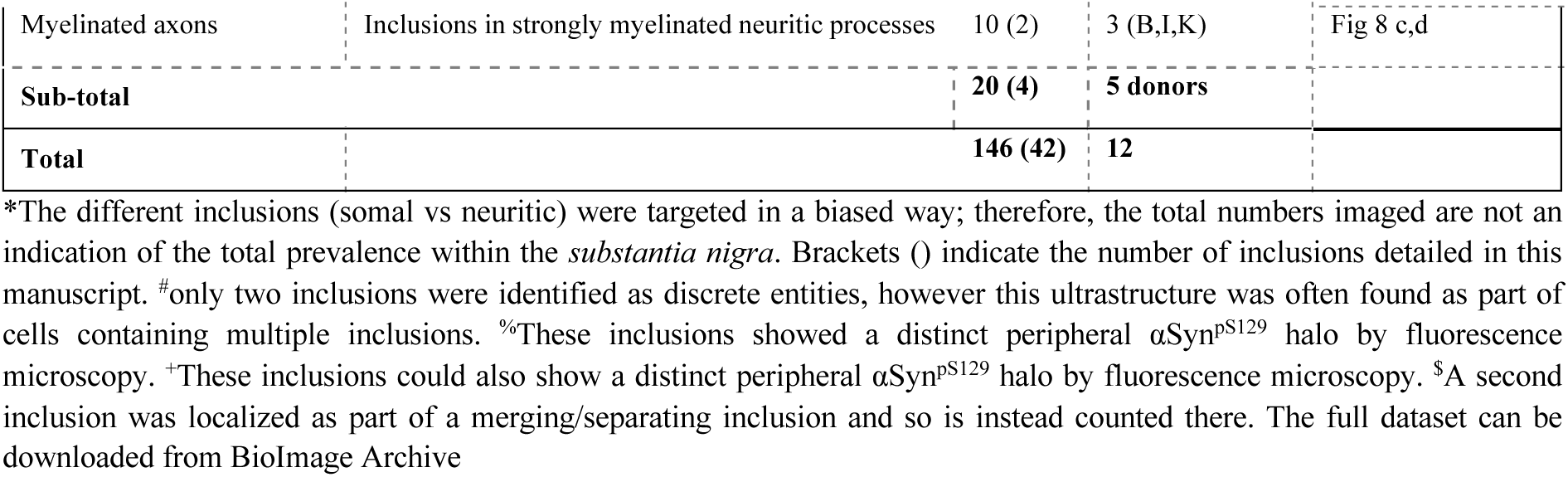
Morphological classification of αSyn^pS129^ inclusions localized by CLEM.

### αSyn^pS129^ pathology in the soma of nigral dopaminergic neurons is distinctly fibrillar

We first characterized the ultrastructure of αSyn^pS129^ inclusions which were clearly inside the soma of nigral dopaminergic neurons (n=59). All somatic inclusions showed a consistently fibrillar ultrastructure, within which several distinct ultrastructural phenotypes were identified, as described below:

#### Diffuse/punctate

Diffuse/punctate αSyn^pS129^ immunoreactivity was often seen interspersed amongst the neuromelanin granules within the neuron. Due to the sparsity of the fluorescent signal in single z-planes throughout the cell, exact correlation of the αSyn^pS129^ immunopositive region with a specific ultrastructure by EM was not possible.

#### Small aggregates

Small aggregation centers (< 5 μm) were localized amongst the neuromelanin granules, and the correlated ultrastructure showed sparse fibrils intermixed with vesicles and proteinaceous material (n=4; Figure 2a). Although it cannot be ruled out that these fibrils belong to the cytoskeleton or other filamentous material, they are likely to be αSyn fibrils since their location corresponds to where the αSyn^pS129^ immuno-fluorescence is brightest. Mitochondria and components of the autophagy-lysosomal pathway were observed around the periphery of the aggregates and sometimes clustered together amongst the fibrils (Figure 2a). The mitochondria close to the aggregates, and further out in the cytoplasm, were often swollen (> 1μm in diameter) and had fragmented cristae, and we also observed a mitophagy event near an aggregate (Figure 2a). Such dysfunctional mitochondrial phenotypes have been previously associated with PD^37,38^.

**Figure 2:**
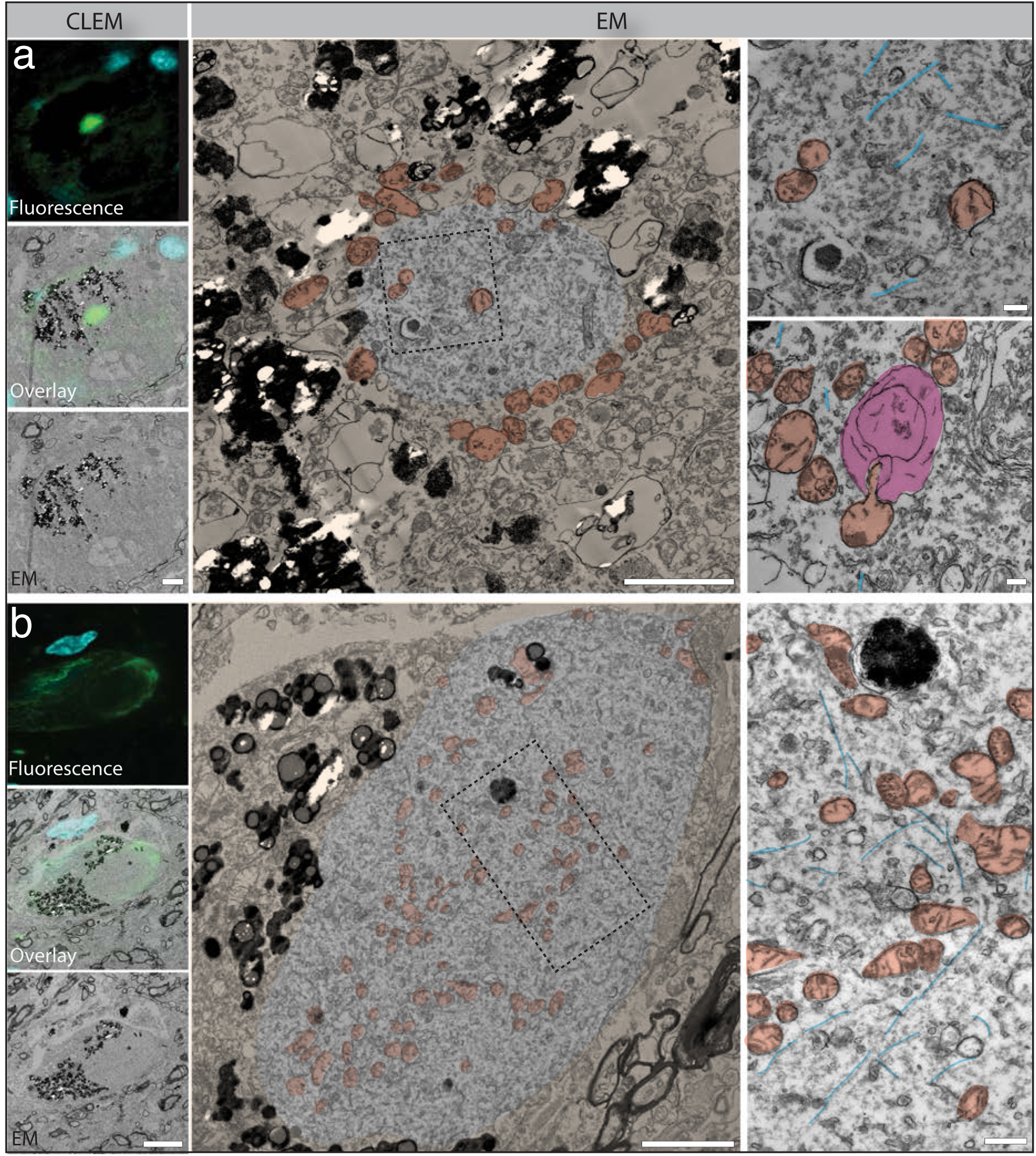
Ultrastructure of small aggregates and pale bodies in neuromelanin cells show dispersed fibrils amongst clustered mitochondria. Inclusions were localized by CLEM^FL^ against αSyn^pS129^ (green; EP1536Y Abcam) and DAPI (cyan). TEM micrographs of a small aggregation center **(a)** and a pale body **(b)** reveal the presence of dysmorphic mitochondria with disrupted cristae (examples coloured in dark orange) clustered together and interspersed amongst sparse fibrillar material (examples coloured in blue). An autophagosome (pink) ingesting a mitochondria near the small aggregation center was observed in an adjacent z-plane. The tissue surrounding the αSyn immunopositive areas are false-coloured light orange for clarity. Scale bars: CLEM - 10 μm; EM - low magnifications 3 μm; high magnifications 500 nm.

#### Large aggregates or pale bodies

Large inclusions (observed up to ∼20 μm diameter) showing diffuse αSyn^pS129^ immuno-fluorescence and distinctly surrounded by neuromelanin granules correlated to an ultrastructure of loosely packed and randomly arranged fibrillar material intermixed with dispersed vesicles and clustered mitochondria (n=13; Figure 2b and Supplementary Fig 1). This ultrastructural arrangement has previously been attributed to pale bodies^10,11^.

#### Halo Lewy bodies

Inclusions showing a distinct peripheral αSyn^pS129^ halo by fluorescence microscopy could be correlated to three distinct fibrillar arrangements by EM with the density of fibril packing assessed by electron tomography and segmentation (Figure 3). The ultrastructures are here classified as: 1-layer inclusions (observed between 3-6 μm diameter), with randomly arranged and loosely packed fibrils (n=2; Figure 3a); 2-layer inclusions (observed between 5-13 μm in diameter), with loosely packed, randomly arranged fibrils at the periphery and a higher density fibrillar mesh at the center (n=17; Figure 3b); and 3-layer inclusions (observed between 9-12 μm in diameter), with loosely packed and linear radiating fibrils on the periphery, a higher density fibrillar mesh in the middle, and an electron-dense central core (n=3; Figure 3c).

**Figure 3:**
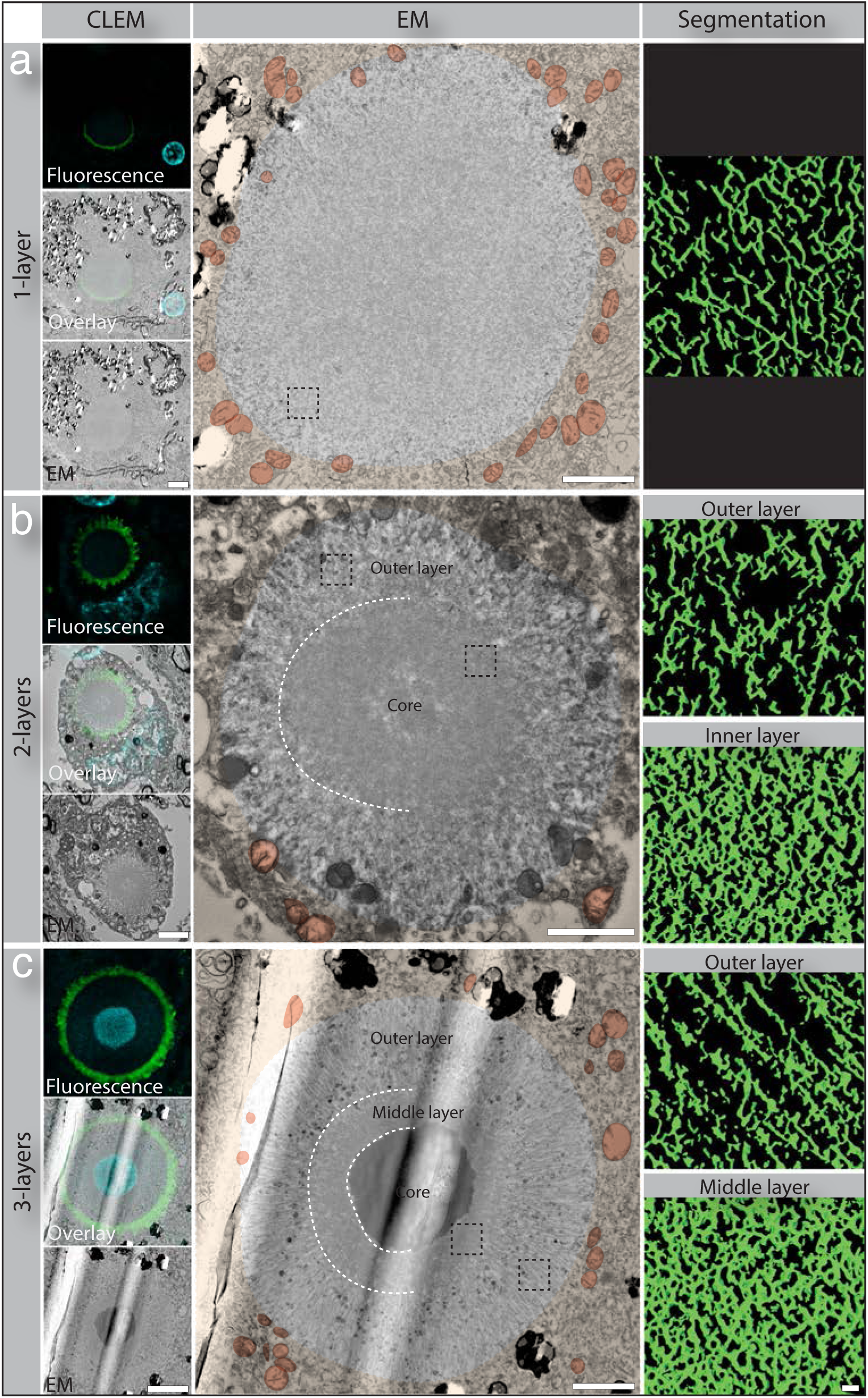
Halo LBs form three distinct ultrastructural arrangements. The inclusions localized within neuromelanin positive neurons by CLEM^FL^ show a halo of αSyn^pS129^ (11a5, Prothena; green). DAPI staining (cyan) is also shown. The EM images show the fibrils within the halo LBs arranged into one, two, or three layers (white dotted lines denote layer boundaries), with mitochondria (examples coloured dark orange) at the periphery of the inclusions. The tissue surrounding the inclusion is false-coloured light orange for clarity. The segmentation of electron tomograms taken from within each layer of the LBs (black dotted square) shows the density and orientation of fibril packing. **(a)** In the one-layer LB, the fibrils are homogeneously distributed and randomly arranged across the whole LB. Also shown in Supplementary Fig. 3 **(b)** In a two-layer LB the outer layer shows a lower density fibril packing compared to the fibrillar mesh that makes up the center. **(c)** The three-layer LB shows a linear arrangement of fibrils in the outer ring, a dense fibrillar mesh in the middle core and a highly electron dense center. A strong DAPI signal was observed in the electron-dense core of the 3-layer LB. The vertical streaks in the EM are due to a tear in the carbon film on the EM grid. Scale bars: CLEM - 4 μm; EM 2 μm; segmentation 50 nm.

The increasing size combined with the increasing level of structural arrangement suggests that these layered ultrastructures represent different stages of LB maturation, starting from a smaller 1-layer LB and ending with the larger 3-layer LB. For all three types, dystrophic mitochondria were consistently observed in the immediate periphery of the fibrillar material, with vesicles of various sizes interspersed amongst the peripheral fibrils. The 2-layer inclusions were observed most frequently (n=17), with the 1-layer (n=2) and 3-layer (n=3) inclusions being observed comparatively rarely.

A clear ring of neurofilament immunostaining could be seen surrounding some halo LBs (Supplementary Figure 2), consistent with previous reports^29^. In contrast to the apparent structural organization implied by the immunostaining, the correlated EM did not reveal a distinct neurofilament ring or a clear ultrastructural boundary between neurofilament and αSyn fibrils (Figure 3). This discrepancy could reflect a limitation of 2D room-temperature TEM in resolving the 3D arrangement of complex filamentous networks and may therefore require volumetric EM for definitive visualization. Additionally, neurofilament may be interspersed within the αSyn fibrillar network and so not visually obvious or present in a form below the resolution of room-temperature EM, such as fragmented or degraded assemblies.

Additionally, DAPI staining was seen in the center of the halo LBs, most prominently in the three-layered LB (Figure 3c and Supplementary Figure 2). Previously reported, this staining has been hypothesized to reflect the presence of nucleic acid material^22,39^. However, due to the exceptionally high electron density of the Lewy body core, where the strongest DAPI signal was observed, it was not possible to resolve the underlying ultrastructure in sufficient detail to determine the molecular identity of the material giving rise to the DAPI signal.

In other examples of halo LBs, one had peripheral radiating fibrils and a dense core at its central slice, similar to a 3-layer LB (Figure 4a). However, the core was not as electron dense as for other 3-layer LBs, looking instead to be composed of a dense fibrillar mesh as observed for 2-layer LBs. Additionally, a third layer between the outer peripheral fibrils and dense core was not as clearly defined as for other examples of 3-layer LBs. Therefore, this inclusion appears to be an intermediate between a 2-layer and 3-layer LB supporting the idea that these “layered” ultrastructures represent different stages of halo LB maturation. In a further example of a 3-layer LB, the electron dense core was not spherical, as has been previously depicted in the literature, but is elongated and irregularly shaped (Figure 4b). With a width measuring 17 μm at its longest point, this inclusion was much larger than the other 3-layer LBs observed and therefore could represent two 3-layer LBs merging. Alternatively, it could imply that even at the mature stages, the 3-layer LBs are not solid structures, but are highly dynamic.

**Figure 4:**
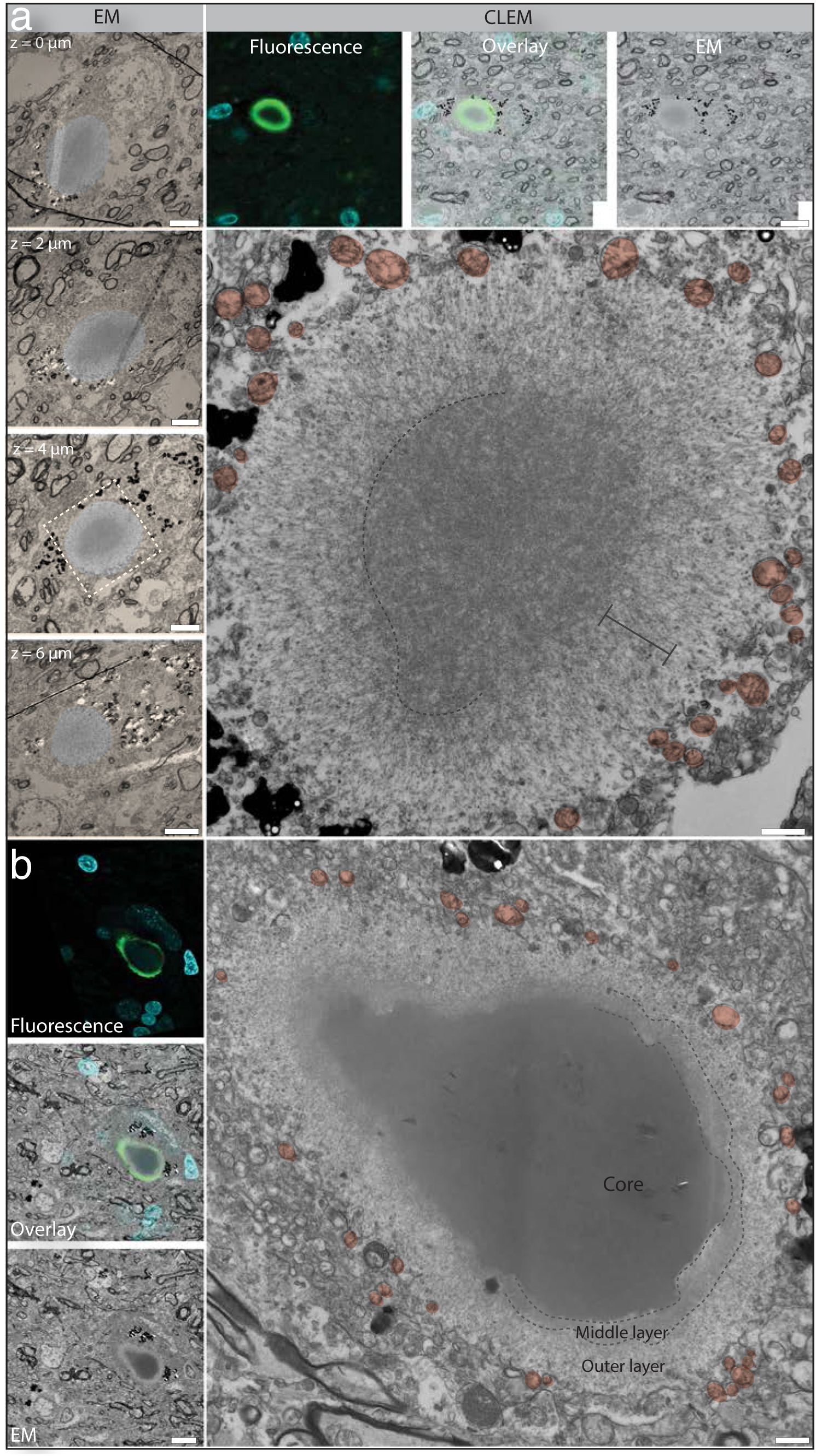
Hybrid halo LB ultrastructures. **(a)** The EM of this inclusion shows linear radiating fibrils in the outer layer similar to a 3-layer LB. The core is less dense than for a typical 3-layer LB, even when examined across different z-planes through the inclusion volume, and is composed of a fibrillar mesh similar to a 2-layer LB. There is no clearly defined middle layer, and there is an increased density between the radiating fibrils and dense core (gray bracket), suggesting this inclusion may be a transition between a 2-layer and 3-layer LB. **(b)** The electron dense core of this 3-layer LB has an elongated and irregular shape, giving the impression of two 3-layer LBs merging together, or suggests that a 3-layer LB can be highly dynamic. Each inclusion was localized by CLEM^FL^ with the αSyn immunostaining (pS129, EP1536Y Abcam; green) and DAPI staining (cyan) shown. The tissue surrounding the inclusions is false-coloured light orange for clarity. Examples of peripheral mitochondria are coloured dark orange. Scale bars: CLEM 10 µm; high magnification EM 1µm.

It is important to note that each of the three ‘layer’ ultrastructures we observed for halo LBs have been previously described in the literature. However, the examples were from different patients, with differences in the (often-long) post-mortem delay, tissue fixation methods and the protocols used for EM preparation. As many of these examples displayed significant ultrastructural extraction in the surrounding tissue ultrastructure, it was not clear if the observed ultrastructural differences were due to preparation or post-mortem artifacts^40^. In contrast, our samples have excellent tissue ultrastructural preservation, and we observed examples of each type of halo LB within the same patient (Supplementary Fig 3a), supporting our hypothesis that these “layered” ultrastructures are different stages of halo LB maturation.

When overlaying the fluorescence and EM for the halo LB’s, we noted that the pattern of αSyn^pS129^ staining by fluorescence did not correlate with the spatial distribution of total fibrillar material in the LB but was instead confined to the peripheral edge of the outermost fibrillar layer (Figure 3). One possible explanation is limited antibody penetration due to the dense packing of fibrils, particularly in the central core of the 3-layer LB, whereas the small-molecule dye DAPI is able to penetrate these regions more readily. Consistent with this, fluorescence microscopy studies have previously reported a laminar organization of proteins and αSyn epitopes within LBs^22,41^.

To address this, we assessed αSyn^pS129^ IHC staining on ultrathin resin sections where the epitopes are accessible across the entire cross-section of the LB. Here the staining pattern matched more precisely with the ultrastructural fibrillar distribution, but still also showed a pronounced peripheral density for αSyn^pS129^ (Supplementary Figure 3b). In contrast, IHC on adjacent sections with an antibody targeting full-length αSyn (SYN1, BD Biosciences) showed homogenous staining across the LB, even in regions of densely packed fibrils. Together these data suggest that some αSyn epitopes remain accessible, despite the increasing density of fibril packing towards the core of a LB, whereas others, such as the pS129 epitope, may exhibit differential accessibility depending on the local fibril density. As such, the peripheral enrichment of αSyn^pS129^ staining may represent a combination of epitope availability and spatial organization, consistent with previous reports^22,41^, but does not conclusively exclude αSyn^pS129^ from the core of the LBs.

We also observed examples of dopaminergic neurons containing multiple halo LBs (n=17; Figure 5, Supplementary Figure 4). The halo LBs were either connected by patches of a pale body type ultrastructure of sparse fibrillar material intermixed with clustered mitochondria (Figure 5a,b), or they showed intermixing of the peripheral fibrils at the boundary between the LBs (Figure 5c,d). In some z-planes the boundary between two LBs could not be easily distinguished giving the distinct impression of organizational fluidity (Figure 5c). The halo LBs were most often either the 1- or 2-layered ultrastructures. In one instance, we observed a multi-halo LB cell which contained all three types of the layered halo LB ultrastructures (Figure 5d). Notably, the 3-layer LB in this cell contained a central core with a much lower electron-density than the other 3-layer LBs observed independently in cells. Taken together, these results support existing data that pale bodies are precursors to halo LBs and that the different layered halo LB ultrastructures are distinct stages of LB formation.

**Figure 5:**
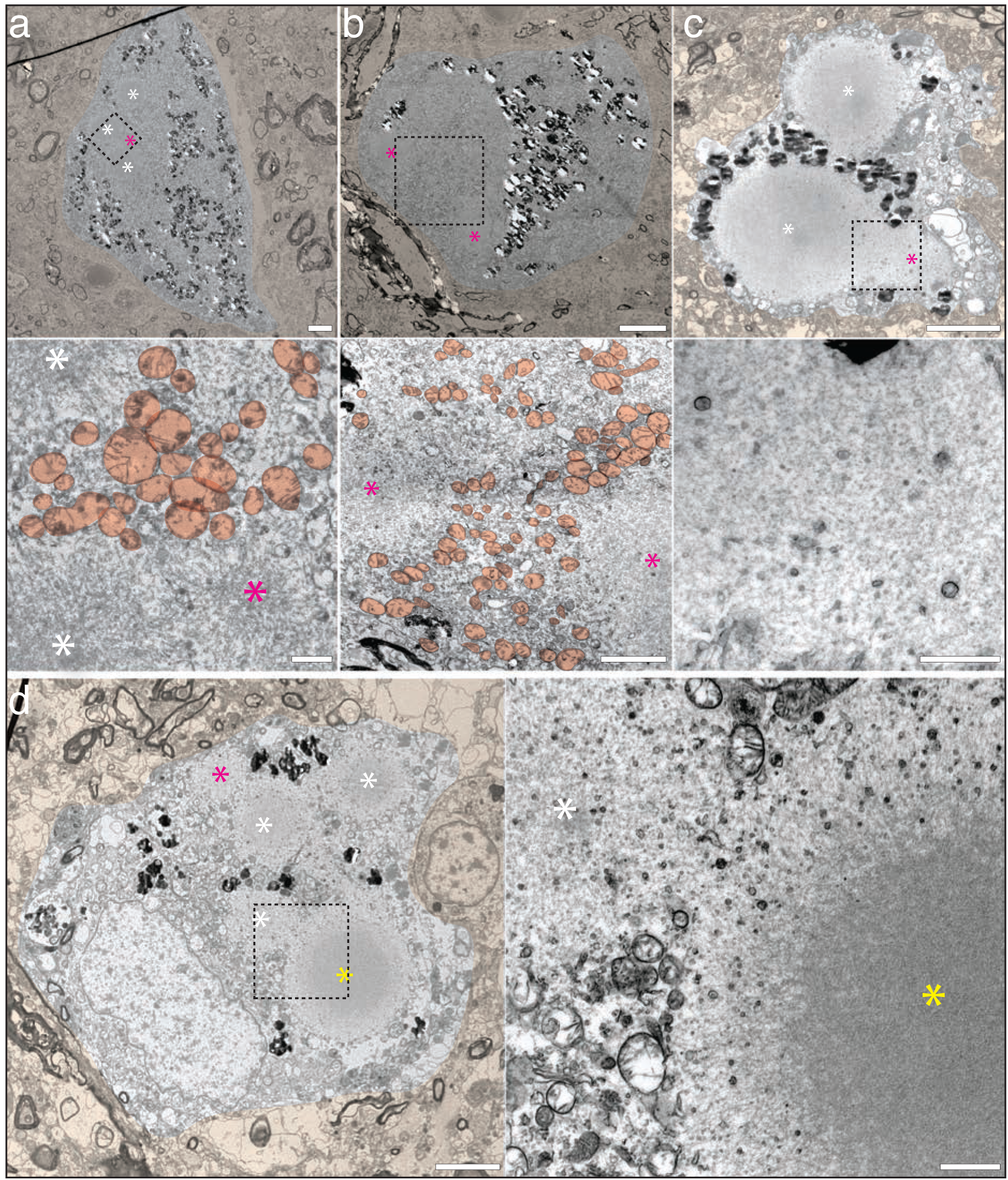
Examples of neurons containing multiple halo LBs. Multiple LBs inside the same neuromelanin containing neuron displaying either the one- (magenta asterisk), two- (white asterisk), or three-layered (yellow asterisk) ultrastructure could be connected by sparse fibrils intermixed with clustered mitochondria (examples coloured orange; **a,b**) or were directly intermixed at their fibrillar boundaries (**c,d**). Areas inside black boxes are shown at higher magnification. The tissue surrounding the cell containing the inclusions is false-coloured light orange for clarity. The CLEM localization for these inclusions is shown in Supplementary Figure 4. Scale bars: low magnification 5 μm; high magnification 1 μm.

Finally, we additionally observed αSyn inclusions (n=2) in neurons exhibiting clear morphological signs of degeneration or pathological stress (Supplementary Figure 5). In these examples, the nuclei appeared shrunken, compacted and electron dense, the cytoplasm, also compacted and electron dense, contained increased vacuoles. Although these vacuoles did not display a clearly defined dense core, they may represent granulovacuolar bodies^42^. Such structures are most commonly described for tau-positive neurons but also thought to reflect a generalized neuronal stress response to intracellular protein aggregation^42,43^. In one example, the fibrillar organization of the αSyn inclusion was that of a 1-layer LB, and the other contained both a 1- and 2-layer LB. In both neurons, the neuromelanin was still clearly visible.

### Compact neuritic αSyn pathology displays ultrastructural heterogeneity that is rich in membranes

We next characterized the ultrastructure of large aggregates (>5 μm) that could not be directly associated to neuronal soma by fluorescence microscopy, as there were no clear cellular demarcations such as an adjacent nucleus, no clear cellular neurofilament staining, or neuromelanin in the immediate vicinity of the aggregate (n=67; Figure 1e). They were also not classified as bulgy Lewy neurites as their dimensions were not elongated in one dimension (XY, or Z), as would be expected for a Lewy neurite (Figure 1f) but were roughly spherical in 3D. As stated previously, the ring of neurofilament staining surrounding the inclusions indicated they were localized within a neuritic compartment and were therefore classified as compact neuritic inclusions.

By EM, we observed ultrastructures for these inclusions rich in densely packed membranous material (Figure 6), as similarly described in our previous study^18^. The spatial distribution of membranes in these inclusions was completely encompassed by the αSyn^pS129^ immunostaining and showed an ultrastructure heavily dominated by vesicular structures and membrane fragments. This composition significantly differed from the mitochondria-rich pale-bodies which only showed sparse vesicles and membrane fragments (Figure 2B and Supplementary Figure 1).

**Figure 6.**
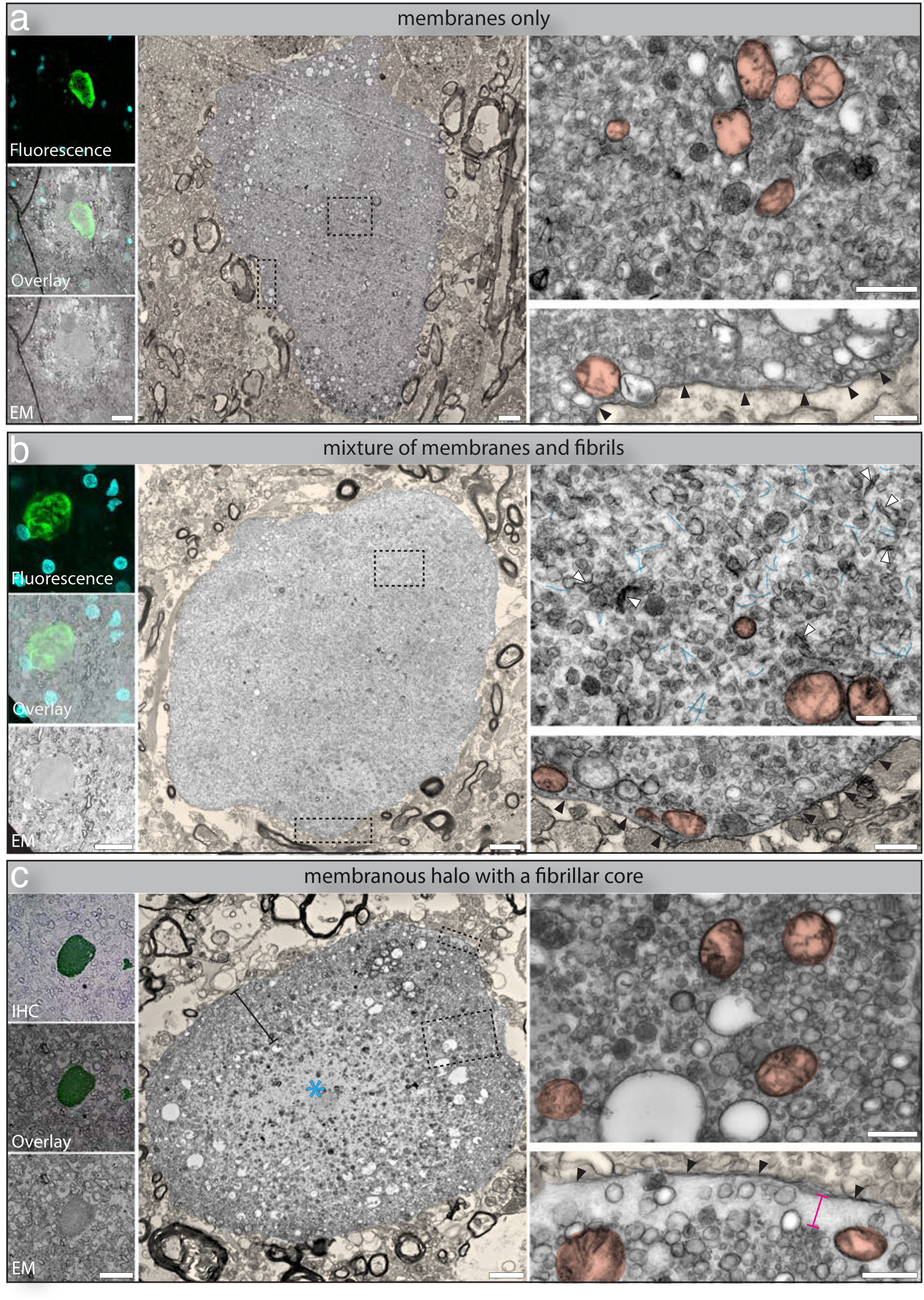
Ultrastructural heterogeneity of compact neuritic pathology. The ultrastructure of inclusions localized by CLEM^FL^ shows (**a**) densely packed membranous material with no detectable fibrillar material or (**b)** densely packed membranous material intermixed with fibrils. αSyn immunostaining (pS129, EP1536Y Abcam; green) and DAPI (cyan) is shown. **(c)** An example of an inclusion localized by CLEM^IHC^ showing a membranous halo (black bracket) surrounding a fibrillar core (blue asterisk). αSyn immunostaining (LB509, Life Technologies; green) overlaid on the toluidine blue image from an adjacent section. Higher magnification EM images from within the inclusions are shown (black dotted squares). Examples of mitochondria are coloured in dark orange, fibrils in blue and membrane fragments denoted with white arrowheads. The surrounding ring of neurofilaments (magenta bracket) and neuritic membrane (black arrowheads) are indicated where visible. The surrounding background tissue has been false-coloured light orange for clarity. Scale bars: CLEM 20μm; low magnification EM 2 μm, high magnification EM 500 nm.

The non-somal inclusions displayed a wide variety of ultrastructural arrangements. They could be composed of densely packed membranous material, devoid of any fibrillar material (n=3; Figure 6a and 7). While these large membranous-only inclusions were observed only twice, additional small (<5 µm) membranous-only aggregates were observed with higher frequency (n=6; Supplementary Fig. 6). In other examples, the membranous material was uniformly distributed and intermixed with fibrils (n=10; Figure 6b) or, showed dense fibril accumulations surrounded by a distinct halo of densely packed membranous material of varying thickness (n=19; Figure 6c, Supplementary Figure 7). Finally, some inclusions were predominantly fibrillar, with no specific ultrastructural arrangement of the fibrils, and a halo of sparsely distributed membranous material located towards the periphery (n=23; Supplementary Figure 8).

**Figure 7.**
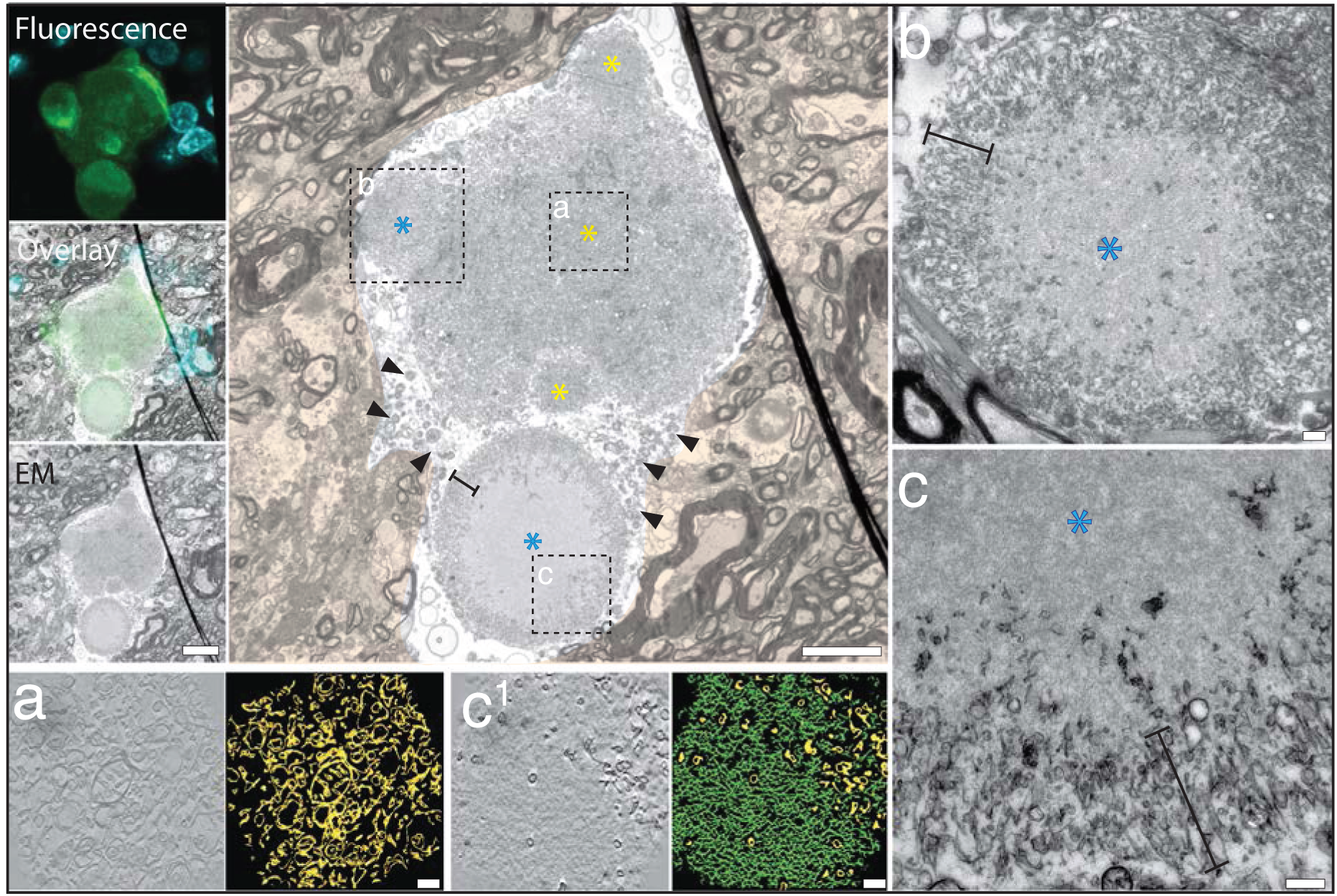
Distinct and multiple ultrastructures within the same neuritic inclusion. An inclusion localized by CLEM^FL^ with αSyn^pS129^ immunostaining (11a5, Prothena; green) and DAPI (cyan) is composed of a main central aggregate and four peripheral puncta. By EM, the main central aggregate plus two peripheral puncta show an ultrastructure of densely packed membranous material (yellow asterisk). The segmentation of an electron tomogram taken from within the central aggregate **(a)** did not reveal the presence of any fibrillar material (also see Supplementary Figure 9). The two other peripheral puncta show an ultrastructure of a membranous halo (black brackets) and fibrillar core (blue asterisk) **(b,c)**. The inclusion shown at higher magnification in (b) is taken from an adjacent z-plane (−1.8 μm) to show the clear membranous halo and fibrillar core. The halo of the larger puncta (c) is composed mainly of membrane fragments. For the z-plane shown, this inclusion is connected to the central aggregate by cellular material such as mitochondria and large vesicles (black arrowheads). The segmentation of an electron tomogram taken from within this inclusion (c^1^) shows the highly dense packing of the fibrillar core (green) contained within the membranous halo (yellow). Scale bars: CLEM 10 μm; low magnification EM 5 μm, high magnification (b,c) 500 nm; tomography and segmentation (a,c^1^) 20 nm.

A defining feature of these neuritic inclusions was the presence of a membrane surrounding the αSyn immunopositive regions (Figure 6, Supplementary Figures 6-8; black arrows), ruling out the possibility of them being extracellular LBs^23^. A recent cell model using iPSC-derived dopaminergic neurons described similarly membrane-rich αSyn inclusions that were membrane-bound, and the authors suggested that LBs may therefore form within the lumen of endo/lysosomal organelles^44^. However, in our observations a ring of organized cytoskeleton filaments (likely corresponding to the ring of neurofilament staining observed by immuno-fluorescence (Figure 1f)) could often be observed between the boundary membrane and the αSyn immunopositive region (Figure 6c, Supplementary Figures 7c and 8b), ruling out the possibility that they are enlarged lysosomes. Additionally, the membrane could sometimes be observed extending along a projection leading away from the αSyn immunopositive area (Supplementary Figures 7b and 8c). Therefore, it is likely that this boundary membrane represents the plasma membrane of the neurite, supporting the idea that these inclusions are compact inclusions within neuritic projections.

**Figure 8.**
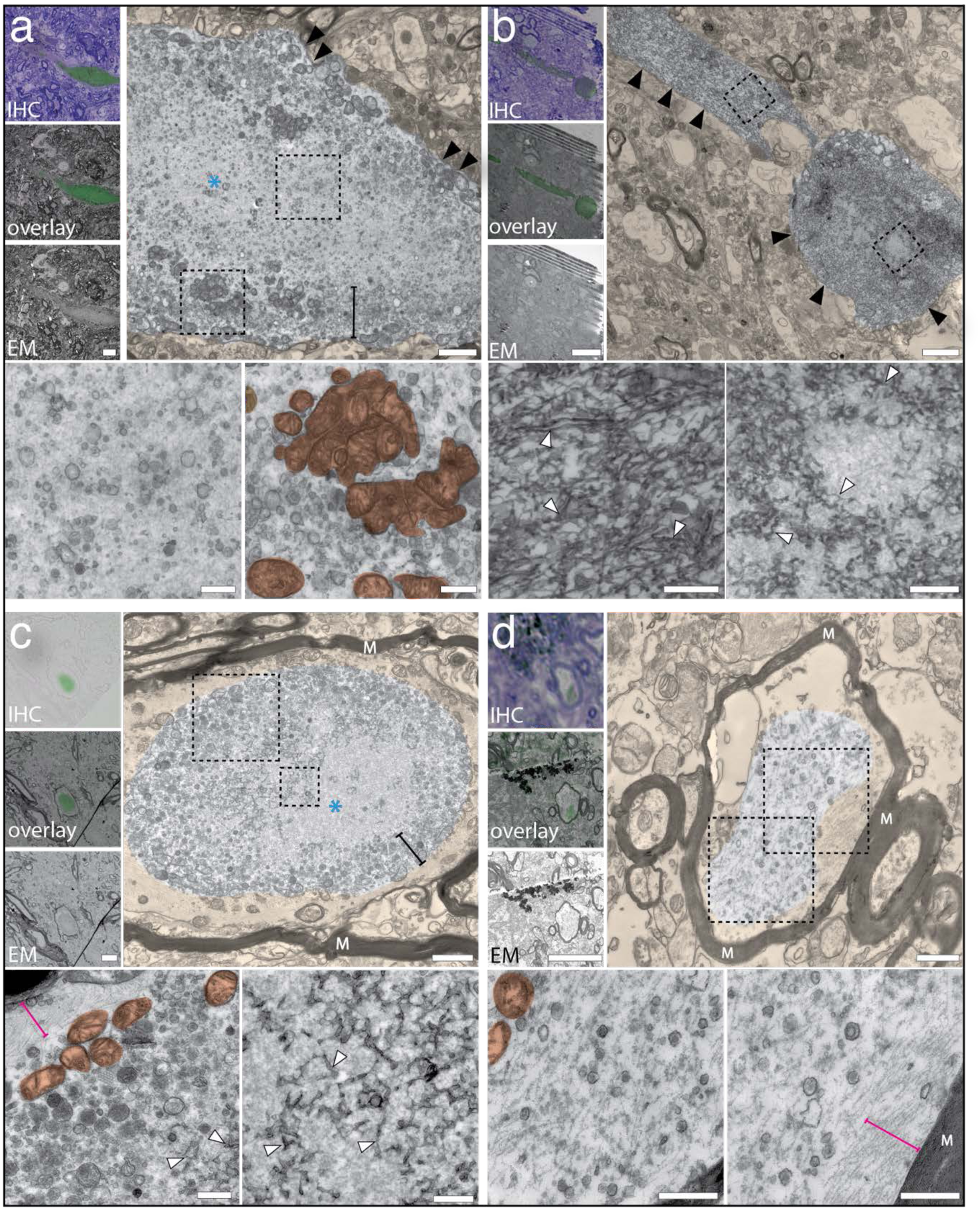
Ultrastructural heterogeneity identified for bulgy Lewy neurites and myelinated axonal αSyn pathology. The CLEM^IHC^ localization showing αSyn immunostaining for bulgy Lewy neurites (a,b) and axonal αSyn pathology (c,d) (a,b: LB509, Life Technologies; c: SYN-1, BD biosciences; d: pS129, EP1536Y Abcam; green) overlaid on the toluidine blue image from an adjacent section. No toluidine blue image was available for panel (c). The ultrastructure of the bulgy neurite shown in **(a)** has a membranous halo (black bracket) surrounding a fibrillar core (blue asterisk), while **(b)** shows fibrils intermixed with membrane fragments. The axonal αSyn pathology shown in **(c)** has an ultrastructure of a membranous halo (black bracket) composed predominantly of vesicles, with some peripheral mitochondria and some internal patches of accumulated membrane fragments surrounding a fibrillar core (blue asterisk). The axonal pathology shown in **(d)** is composed of αSyn fibrils intermixed with vesicles. For all inclusions the myelin sheath (M) or neuritic membrane (black arrowheads) are indicated. Areas indicated by black dotted squares are shown at higher magnification. Mitochondria are coloured in orange, membrane fragments are indicated with white arrowheads and cytoskeleton filaments are indicated by a magenta bracket. In all low magnification EM images, the tissue surrounding the αSyn immunopositive region is false-coloured light orange. Scale bars: CLEM 10 μm; low mag EM (a,b) 1 μm, (c,d) 3 μm; high mag EM 500 nm.

A particular example of note was a compact neuritic inclusion showing multiple discrete puncta composed of various ultrastructural phenotypes within the same aggregate (Figure 7). By EM, this inclusion consisted of a large central aggregate composed of a dense accumulation of membrane fragments, vesicles and mitochondria, where no fibrillar material could be observed (Figure 7a). Two puncta located on the periphery of the large central aggregate were similarly composed only of densely accumulated membranous material, while a third peripheral puncta showed a membranous halo, fibrillar core ultrastructure (Figure 7b). Two of the three small peripheral inclusions are visually distinct from the main central aggregate due to a difference in density of the surrounding membranous material, with the third being visually separated due to a distinct invagination of the main aggregate.

A fourth larger aggregate was composed of a distinctive halo of membrane fragments surrounding a densely packed and disorganized fibrillar core (Figure 7c,d). In the section shown, this fourth aggregate shares no direct intermixing of boundaries with the main inclusion body, but the two structures are connected by a continuous tract of mitochondria and large vesicles in the surrounding space (Figure 7, black arrows) and show a closer spatial arrangement when observed in adjacent sections (Supplementary Figure 9).

For the main aggregate body, tomography and segmentation additionally did not identify any fibrillar material mixed amongst the densely accumulated membranes (Figure 7a), and neither could any fibrillar material be identified in any of the adjacent sections by EM (Supplementary Figure 9). No nucleus or neuromelanin was observed within the inclusion boundary across the ∼20 um volume for which this inclusion could be followed by EM. Additionally, in one adjacent section, the inclusion appears to be within a projection (Supplementary Figure 9). Collectively, these features support the classification of this multi-punctum aggregate as a compact neuritic inclusion.

The distinctive arrangement of the multiple puncta within this single inclusion gives the impression that they have formed at the periphery of the central membranous aggregate and are in a dynamic state of merging with or separating from the larger aggregate. Similar impressions of merging/separating inclusions were seen in other neuritic inclusions (n=7; Supplementary Figure 10).

### Bulgy Lewy neurites and axonal pathology also show heterogeneous and membrane-rich ultrastructures

We additionally localized several bulgy Lewy neurites (n=10) and αSyn immunopositive inclusions inside myelinated axons (n=10) using CLEM^IHC^ (Figure 8). The bulgy neurites were distinguished by their enlarged, elongated projection through the tissue, where their length was greater than their width (Figure 8 a,b). Axonal pathology was identified by EM from its obvious containment within myelin sheaths (Figure 8 c,d). We found these pathologies also showed the same ultrastructural heterogeneity as seen for the non-somal inclusions; including, membranes intermixed with fibrils, and the membranous halo, fibrillar core ultrastructure. The membrane-rich ultrastructures were dominated by vesicles and membrane fragments, and some clustered mitochondria could also be observed. Peripheral cytoskeleton filaments were observed in immuno-negative areas of the axonal inclusions and were easily distinguishable from the αSyn fibrils due to their highly linear and organized spatial arrangement compared to the densely packed (Figure 8c) or disorganized arrangement of the αSyn fibrils (Figure 8d). The neuritic membrane of the bulgy neurite was clearly visible and, for the axonal pathology, cytoskeleton filaments were observed to separate the αSyn immunopositive region from the myelin sheaths. There was no additional membrane specifically containing the αSyn immunopositive regions for either the bulgy neurites or the axonal pathology, further supporting the idea that the membrane surrounding the compact neuritic aggregates derives from the plasma membrane of the neurite.

## Discussion

Over the past decades, several efforts have been made to describe the staging of LB formation based on the morphological heterogeneity of αSyn immunopositive aggregates observed by IHC or immunofluorescence in human post-mortem brain tissue ^22,32–34^. In this study, we used CLEM to characterize the ultrastructure of αSyn^pS129^ aggregates in the *substantia nigra* and found that αSyn^pS129^ pathology could be categorized into two distinct ultrastructural subtypes according to their subcellular localization: fibrillar inclusions within the soma of dopaminergic neurons and membrane-rich inclusions within neuronal processes.

Inclusions located within the soma of dopaminergic neurons in the *substantia nigra* consistently exhibited fibrillar-only ultrastructures, in agreement with previous ultrastructural studies associating fibrillar αSyn with classical halo LBs^6,45^. Across all 12 donors examined, the somal inclusions ranged from small punctate cellular staining to the large classical halo LBs, displaying increasing size, fibril density, organization into layered morphologies and characteristic redistribution of mitochondria. Taken together these observations support established histological staging models of somal LB maturation^22,32–34^.

Within this framework, our data are compatible with a model in which initially sparse fibrillar material gradually accumulates together with damaged mitochondria to form large pale bodies and subsequently layered halo LBs, with mitochondria progressively excluded towards the periphery of the inclusion (Figure 9). Support for this sequence is provided by our observations of intermediate or “hybrid” ultrastructures and by neurons containing multiple LBs at different apparent stages of organization (Figure 4a, 5; Supplementary Figure 4).

**Figure 9.**
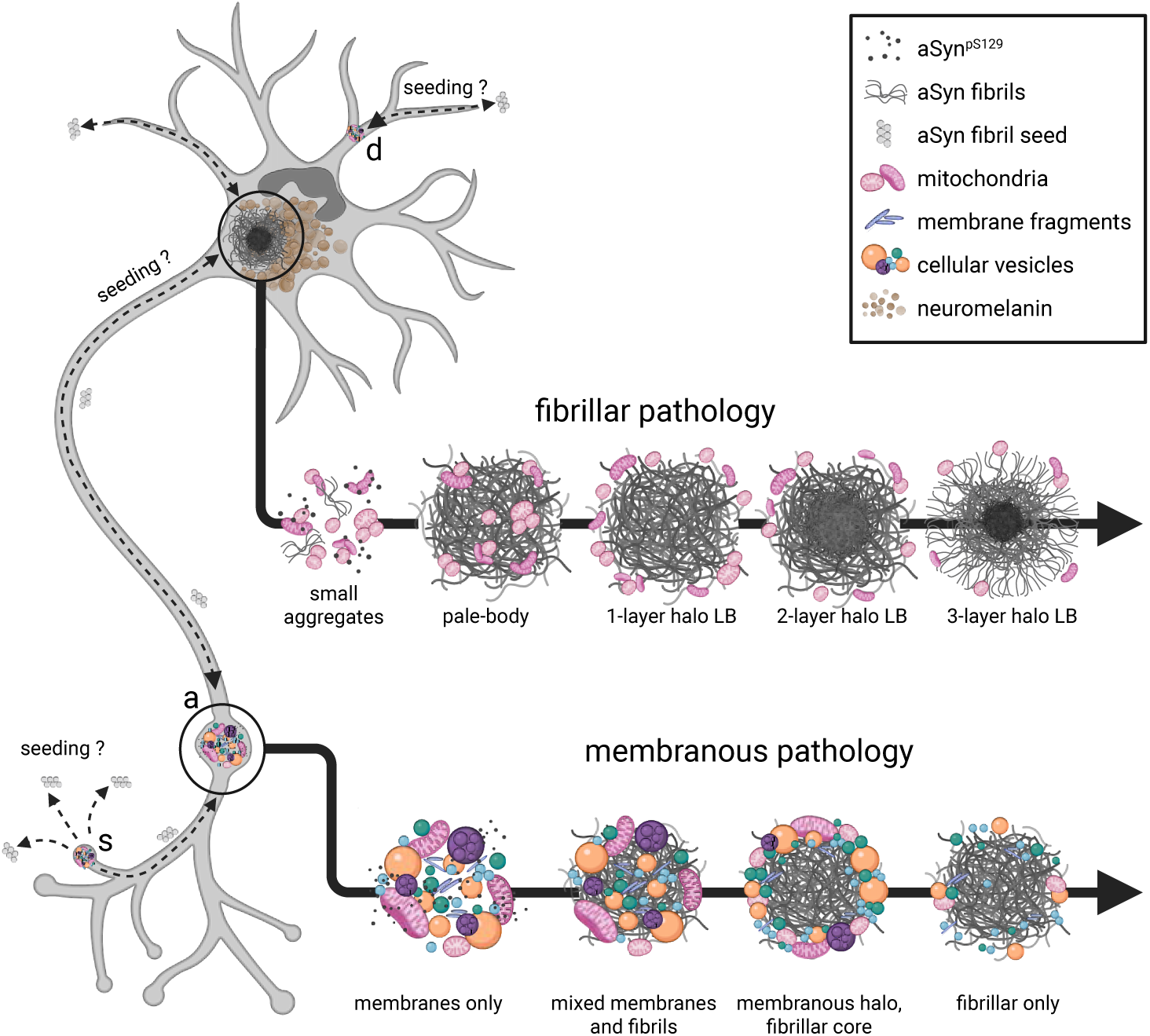
Proposed model for the formation and diversification of fibrillar and membranous αSyn^pS129^ inclusions in PD. Schematic illustrating a working model derived from ultrastructural observations in the *substantia nigra* of post-mortem human brain. Fibrillar αSyn accumulates within the soma of dopaminergic neurons, together with clustered dystrophic mitochondria, forming pale bodies that mature into layered halo Lewy bodies with peripheral mitochondria. In contrast, within neuronal processes (including axons (a), dendrites (d) and potentially synaptic compartments (s)) pathological interactions between αSyn and cellular membranes may give rise to membrane-rich inclusions lacking fibrillar material. These membranous inclusions, characterized by a crowded lipid environment and high local concentrations of non-fibrillar αSyn, may provide a site conducive to αSyn fibril nucleation. Progressive accumulation of fibrillar material could then lead to mixed membrane–fibril assemblies, inclusions with a membranous halo surrounding a fibrillar core, and fibrillar-only inclusions. Whether these two inclusion types arise through independent aggregation pathways or are mechanistically linked (for example through the generation and spread of transmissible αSyn species) remains unresolved. Dashed arrows indicate hypothesized processes that are not directly supported by the present data. This figure was created in BioRender. Lewis, A (2026) https://BioRender.com/vc41699.

In contrast to somal inclusions, αSyn^pS129^ pathology outside the soma was dominated by membrane-rich accumulations displaying marked ultrastructural heterogeneity. These inclusions ranged from densely packed membranous accumulations lacking detectable fibrils to mixed membranous–fibrillar assemblies and structures composed of fibrillar cores surrounded by membranous halos. These inclusions were typically enclosed by a surrounding membrane and frequently associated with a ring of neurofilament (immunostaining (Figure 1f), or ultrastructure (Figure 6 b,d), supporting their classification as compact αSyn^pS129^ aggregates located in neuritic projections. The large diameter (> 5 μm) of these inclusions is likely the result of extensive neuritic swelling, caused by aggregate formation. This interpretation is supported by the presence of similar membrane-rich ultrastructures in bulgy Lewy neurites and axonal αSyn pathology (Figure 8).

Due to the limited thickness of the tissue sections used for CLEM, the precise anatomical position of compact neuritic inclusions could not be resolved. Nevertheless, the frequent accumulation of vesicles within these inclusions could indicate proximity to a synapse or a synaptic origin. This interpretation is consistent with recent human post-mortem evidence demonstrating enrichment of αSyn^pS129^ at dopaminergic synapses in early PD, supporting the idea that synaptic and neuritic compartments represent early sites of pathological αSyn accumulation^46^. Further characterization of the membranous components, combined with higher-resolution spatial imaging approaches, will be required to determine their precise cellular context and origin.

Our observation of multiple membranous ultrastructural states within single compact neuritic inclusions mirrors the transition of pale bodies to halo LBs in the soma and suggests they may likewise encompass multiple stages or parallel aggregation states. Analogous to our model for somal LBs, in which increasing fibril density reflects progressive LB formation, we propose that a similar process may occur in neurites. In this scenario, aggregated membranous material containing a high local concentration of non-fibrillar αSyn^pS129^ could provide an environment that triggers αSyn fibril formation, with fibrils subsequently accumulating in density to give rise to the various membrane–fibril ultrastructures (Figure 9).

Large membranous-only inclusions lacking any detectable fibrillar material were only observed in 3 out of 67 compact neuritic inclusions characterized by CLEM. Despite their limited representation, they were observed across multiple donors (including our previous study where such inclusions were first reported^18^), together with the numerous small membranous-only aggregates (< 5 μm), supporting their relevance as a genuine, pathological state. Their underrepresentation at the ultrastrucural level likely reflects several factors, including a transient phase of pathology, the end-stage nature of our cohort, or an antibody bias associated with the pS129-focused targeting strategy. Other aSyn antibodies may reveal a higher abundance of non-fibrillar aggregates, and, if the membranous-only ultrastructures represent an initial stage of neuritic involvement, they may be more prevalent in earlier disease stages. Importantly, rarity alone does not preclude biological significance, particularly given that other LB morphologies widely regarded as mature end-stage forms were also rarely observed in this cohort.

Notably, membrane-rich ultrastructures were not observed within somal inclusions, indicating a clear subcellular segregation between fibrillar and membranous inclusions. Early ultrastructural descriptions by Forno and Norville of “Lewy body–related swellings” within neuronal processes^5^ closely resemble the compact neuritic inclusions characterized here, suggesting that such membrane-rich pathology has long been observed but not clearly distinguished from classical somal Lewy bodies. The distinct spatial separation of the membranous neuritic and fibrillar somal inclusions suggests that they arise through distinct cellular processes although how, or if, these processes are mechanistically linked remains unclear.

Our previous finding that the majority of Lewy pathology consists of a crowded membranous environment introduced a lipid-centric hypothesis for the formation of LBs^18^ challenging the traditional protein-centric views for PD pathology^6,47–50^. The current data extend this model, suggesting that pathology formation in PD is likely to involve a combination of both membrane- and protein-driven processes. The accumulation of damaged membranes may play an important role in promoting the nucleation of αSyn fibril formation consistent with a recent post-mortem study linking membrane damage to aSyn aggregation^51^, as well as a recently published human iPSC-derived dopaminergic neuron seeding model in which aSyn preformed fibrils were shown to induce membrane damage, which was associated with enhanced aSyn fibril formation^52^.

Consistent with this, we observe neuritic membranous inclusions without the presence of fibrillar material, indicating that membrane aggregation can occur in the absence of mature αSyn fibrils. Within the context of the current prion-like hypothesis for somal LB formation (reviewed in^53^), it is therefore plausible that transmittable αSyn species may be generated within these membranous inclusions and subsequently spread along the neurites, or between neurons, contributing to disease pathogenesis (Figure 9).

Such a process is supported by recent observations in an iPSC model by Lam et al., showing that seeded αSyn^pS129^ inclusions are biochemically and morphologically distinct from those that form spontaneously^54^. Furthermore, the idea that membranous neuritic inclusions precede the classic fibrillar accumulations in the soma of dopaminergic neurons, aligns with other post-mortem studies suggesting that αSyn pathology starts in neuritic compartments^8,46^, as well as various cellular seeding models in which neuritic aggregates precede the formation of somatic inclusions and exhibit distinct morphological and biochemical features^52,55^.

Our proposed sequence represents a model inferred from cross-sectional observations of late-stage post-mortem tissue and does not constitute a directly observed temporal progression. It is also important to note that the proposed transmittable species of αSyn, being oligomers or protofibrils (reviewed in^56^), are below the resolution limits of our methods. Additionally, the end-stage nature of our cohort may lead to an underrepresentation of short-lived or early pathological states, particularly given that post-mortem delay and tissue processing may lead to the loss or dissolution of transient, metastable αSyn assemblies. Consequently, static post-mortem observations cannot establish whether distinct ultrastructural phenotypes arise sequentially or independently. In this context, complementary cellular and seeding-based models capable of resolving temporal progression have demonstrated the emergence of distinct inclusion subtypes and that Lewy body formation is a dynamic, multistep process involving both fibrillization and organelle engagement^52,54,55^. Consequently, whether membranous and fibrillar inclusions form through a defined maturation sequence or represent parallel aggregation pathways remains to be established in vitro, in non-seeding model systems, and through future studies examining the ultrastructural diversity and prevalence of these potential maturation states across different disease stages.

In this study, all inclusions identified by CLEM^FL^ used an antibody against αSyn^pS129^, which labeled both fibrillar and membranous inclusions. Consequently, discrimination between these ultrastructural subtypes was based on their EM morphology and their location in or outside the soma of dopaminergic neurons. With numerous microscopy studies reporting differential αSyn epitope staining patterns across synuclein disease pathologies^22–29^, further CLEM studies incorporating antibodies against additional epitopes may reveal further ultrastructural diversity and refine the distinction between fibrillar and membranous ultrastructures.

Additionally, the incorporation of proximity ligation assays or enhanced αSyn staining to enable the detection of oligomeric or lower molecular weight αSyn aggregates^57–60^ may complement our ultrastructural classification and help refine our proposed staging of Lewy pathology. While this study focused on αSyn^pS129^ immunopositive inclusions within the *substantia nigra* of PD patients, future work examining additional αSyn epitopes, brain regions, and other synucleinopathies will be essential to fully capture the molecular and ultrastructural heterogeneity underlying PD pathology.

In conclusion, our study marks the most comprehensive ultrastructural exploration to date, capturing the wide morphological spectrum of Lewy pathology within the *substantia nigra* of PD patients. Our findings delineate membranous and fibrillar αSyn inclusions as discrete entities, suggesting they may arise through distinct aggregation processes, although how and if these pathways are mechanistically linked remains to be determined.

Further mechanistic investigations are now required to elucidate the initial aggregation mechanism of αSyn in association with membranous material, to define the cellular origin of the accumulated membranes, and to determine the specific structural state of αSyn in these inclusions.

Together, our study provides insights into the ultrastructural complexity of αSyn^pS129^ pathology in the human brain and further highlights a potential role for membrane interactions in driving αSyn aggregation. A deeper understanding of these processes may prove critical for developing future therapeutic interventions aimed at modifying disease onset and progression.

## Methods

All donors participated in the brain donation program from the from the Normal Ageing Brain Collection Amsterdam (NABCA: www.nabca.eu) and the Netherlands Brain Bank (NBB: www.brainbank.nl) and provided written informed consent for a brain autopsy and the use of the material and clinical information for research purposes in accordance with the Declaration of Helsinki (www.wma.net/policies-post/wma-declaration-of-helsinki/). Detailed neuropathological and clinical information was made available, in compliance with local ethical and legal guidelines, and all protocols were approved by Amsterdam UMC institutional ethics review board (NABCA 2018/150; NBB 2019/148). The use of human post-mortem tissue at EPFL (Lausanne, Switzerland) was approved by the Cantonal Ethics Committee of Vaud (CER-VD; approval numbers 2020-01692 and 2025-01061), in accordance with Swiss Human Research Act guidelines.

### Human post-mortem brain samples

We included 12 brain donors (Donors A-L) from patients who clinically presented with PD or PD with Dementia (PDD) with a post-mortem delay of <7 hrs (Supplementary Table 1). Demographic features and clinical symptoms were abstracted from the clinical files, including sex, age at symptom onset, age at death, disease duration, presence of dementia, core and supportive clinical features for PD or PDD.

The brain was dissected at autopsy according to the protocol of the NBB (https://www.brainbank.nl/brain-tissue/autopsy/). For electron microscopy, the mesencephalon was collected at the level of the oculomotor nerve and subsequently the *substantia nigra* was dissected into 0.5 cm^3^ tissue blocks at autopsy using a standardized procedure by a neuroanatomist (WvdB). These tissue blocks from the *substantia nigra* were immersion fixed in 4 % paraformaldehyde and 0.1 % glutaraldehyde in 0.15 M cacodylate buffer, supplemented with 2 mM calcium chloride, pH 7.4 for 24 hours and stored afterwards in 0.1 paraformaldehyde in 0.15 M cacodylate buffer until further processing for EM.

For pathological diagnosis, six or seven µm-thick FFPE-embedded sections were immuno-stained using antibodies against αSyn (clone KM51, 1:500, Monosan Xtra, The Netherlands), amyloid-β (clone 4G8, 1:8000, Biolegend, USA) and phosphorylated tau (p-tau, clone AT8, 1:500, Thermo Fisher Scientific, USA) or processed for Gallyas silver staining, as previously described). Braak αSyn stages were determined using the BrainNet Europe (BNE) staging criteria^32^. Based on Thal phases^61^ (amyloid-β), Braak neurofibrillary stages^62^ (AT-8 phospho-tau) and CERAD neuritic plaque scores^63^ (Gallyas method), levels of AD pathology were determined according to NIA-AA consensus criteria^64^. Additionally, Thal CAA stages^65^, presence of aging-related tau astrogliopathy^66^ (ARTAG), and microvascular lesions and hippocampal sclerosis were assessed.

### CLEM

For the fluorescent microscopy, 40-60 μm free-floating brain sections were prepared with a vibratome (Leica VT1200) and immuno-labeled with primary antibodies (αSyn: 11a5 – Prothena - 1/1000, or αSyn^pS129^ EP1536Y - Abcam # 51253 - 1/1000; neurofilament: Sigma Ab5539 - 1/500) overnight at 4 °C and visualized with Alexa-conjugated secondary antibodies (donkey 488 Molecular Probes A32787 - 1/400, chicken 647 Molecular Probes A32933 - 1/400) and DAPI (Biolegend #422801 - 1/800 dilution) to label cell nuclei after incubation at room-temperature for 30 mins. 3D fluorescent images were acquired on a confocal TCS SP8 or a Thunder Tissue imager (Leica Microsystems) using a 63 x oil objective (HC PL APO/1.4 numerical aperture). The sections were then resin embedded using a tissue autoprocessor (Leica Microsystems) with the following incubations: 2% reduced osmium tetroxide for 60 mins, 1% thiocarbohydrazide for 20 mins, 2% osmium tetroxide for 60 mins, 2% uranyl acetate for 60 mins, lead aspartate (pH 5.5) for 40 mins at 60 °C, acetone dehydration (30%, 70%, 100% x 3) over 4.5 hours and epon infiltration (30%, 70%, 100% x 4 over 20 hours. The detailed step-by-step protocol is available at Shafiei et al^36^. Trapezoid blocks containing the αSyn pathologies were removed from the resin-embedded section using laser capture microdissection (Leica LMD7) and the resulting blocks glued on the surface of a resin stub. Serial sections of 90 nm (for EM) and 200 nm (for LM) were cut using an ultracut UC7 (Leica Microsystems) and collected consecutively on EM grids and glass slides, respectively.

Immunohistochemistry was performed on the glass slides. The sections were etched in a saturated potassium ethoxide solution for 3 mins followed by washing in PBS. Endogenous peroxidases were quenched with 1 % hydrogen peroxide in 10 % methanol, before blocking in Dako REAL antibody diluent (Agilent). Antigen retrieval and primary incubations were carried out as outlined in Supplementary Table 2, before washing in PBS supplemented with 0.25 % Triton X and incubation in secondary antibody (ImmPRESS Reagent Anti-Mouse Ig, or Anti-Rabbit Ig, Vector Laboratories) for 30 mins at room temperature. Bound antibody complexes were detected using the permanent HRP Green Kit (Zytomed Systems) with incubation for 3 mins at room temperature. Sections were counterstained with hematoxylin, dehydrated and mounted under glass coverslips for imaging. Adjacent slides were stained with 1% toluidine blue solution. Light microscopy images of selected glass slides containing αSyn immunopositive structures were collected using a Thunder Tissue imager microscope equipped with a DMI8 color camera (Leica).

TEM images were collected at room temperature on a 120 kV Tecnai G2 Spirit TEM microscope operated at 80 kV with a LaB6 filament and a side mounted EMSIS Veleta camera, a CM100 Biotwin (Philips) operated at 100 kV or a Tecnai Spirit BioTwin (FEI) operated at 80 kV with Lab6 filaments and bottom mounted TVIPS F416 cameras.

### Tomography

Tomograms were collected with a pixel size of ∼0.5 nm on a Jeol 2100 Plus at 200 kV equipped with a LaB6 filament and TVIPS camera, or a Talos F200C (ThermoFisher Scientific) operating at 200 kV equipped with an X-FEG electron source and a Ceta camera. Exposures of 0.5 seconds were collected every 2 degrees from −60 to +60 degrees. Tomograms were reconstructed from recorded tilt series using IMOD^67^, binned by a factor of 2 and filtered using a non-local means filter in Amira version 2021.2 (ThermoFisher Scientific). Segmentation of the tomograms was carried out using the semi-automated convolutional neural network protocol of EMAN2^68^ and refined using the UCSF Chimera package^69^ and Amira. The final segmentations in the figures show the central 15 slices of the total segmented volume.

### Image adjustment

Fluorescent images were deconvoluted using the Leica Thunder algorithm (Leica microsystems, Germany). Brightfield microscopy images were corrected for color-blind readers by replacing the red channel with magenta using FIJI^70^. Both brightfield and EM images were adjusted for brightness and contrast where necessary. EM and fluorescent overlays were created in Adobe Photoshop (2024) using the ‘screen’ transparency option for the fluorescent image. Toluidine blue/EM and IHC overlays were made in Adobe Illustrator by ‘multiplying’ the IHC image with the Toluidine blue/EM.

### Statistics and Reproducibility

This study is based on targeted correlative light and electron microscopy (CLEM) analysis of human postmortem brain tissue and was not designed as a repeated experimental assay. Instead, reproducibility is supported by the consistent observation of ultrastructural features across multiple independent human donors and inclusions. In total, 143 αSynpS129-immunopositive inclusions were identified and characterized from the substantia nigra of 12 Parkinson’s disease donors. All inclusions were classified based on subcellular localization and ultrastructural features, and the distribution of morphological classes across donors is summarized in Table 1. Donor information is provided in Supplementary Table 1. Representative micrographs shown in the figures were selected from this larger dataset to illustrate the range of observed morphologies. In total, 40 inclusions are presented in detail in this study. The full dataset is available via the BioImage Archive. No statistical methods were used to predetermine sample size.

## Data Availability

The raw EM micrographs from the full dataset generated in this study have been deposited in the BioImageArchive (http://www.ebi.ac.uk/bioimage-archive) under accession number S-BIAD3276. Additional data is available upon request.

## Acknowledgements

We are grateful to the individuals who participated in the brain donation program of the Netherlands Brain bank and their families, making this study possible. We would like to thank Annemieke Rozemuller and the members of the Netherlands Brain Bank autopsy team for facilitating the collection of high-quality post-mortem brain tissue for EM. We thank the staff at the EM Facility at the University of Lausanne, and Inayathulla Mohammed and Julika Radecke from the Laboratory for Biological Electron Microscopy at EPFL for sample preparation and/or TEM assistance and maintenance, Prothena for providing the pS129 11A5 antibody and the Advanced Optical Microscopy Core O|2 (www.ao2m.amsterdam) for support with confocal imaging.

## Author contributions

AJL, WVdB and HS designed the study. WVdB performed rapid autopsies of PD brain donors and controls, collected brain tissue for EM and performed neuropathological assessment. AJL, LvdH, MDF, NS and DB performed CLEM with assistance from KB, DP, KE and SO. EH and JGJMB assisted with the multi-labelling immuno-fluorescence and fluorescence microscopy. AJL performed the tomography and segmentation. AJL and LvdH prepared the figures. AJL wrote the manuscript. All authors contributed to the analysis and interpretation of the data and approved the final version of the manuscript.

