## Supplementary information for "Ultrastructural diversity and subcellular organization of nigral Lewy pathology in Parkinson’s disease"

**Supplementary Table 1. Demographic neuropathological details of the included PD brain donors used for this study.**

| ID donor | age at death (y) | F/M | ABC | Braak aSyn stage | Braak NFT stage | Thal Phase | CAA type | Number of inclusions by CLEM* | |
|---|---|---|---|---|---|---|---|---|---|
| | | | | | | | | Somal | Neuritic |
| A | 71 | M | A1B0C0 | 6 | 1 | 1 | 0 | 9 | 17 |
| B | 88 | M | A2B2C2 | 6 | 3 | 3 | 2 | 24 | 18 |
| C | 85 | M | A1B1C0 | 6 | 2 | 0 | 0 | 3 | 1 |
| D | 83 | M | A1B2C0 | 6 | 3 | 1 | 0 | 3 | 4 |
| E | 87 | F | A1B2C0 | 6 | 3 | 1 | 0 | 2 | 6 |
| F | 63 | M | A1B1C0 | 6 | 1 | 1 | 0 | 2 | 12 |
| G[#] | 90 | F | A2B2C2 | 6 | 3 | 3 | 2 | 0 | 1 |
| H | 74 | M | A3B2C1 | 6 | 4 | 5 | 1 | 2 | 4 |
| I | 84 | M | A0B1C0 | 5 | 2 | 0 | 0 | 0 | 6 |
| J | 74 | M | A2B1C1 | 6 | 2 | 3 | 1 | 2 | 5 |
| K | 75 | M | A2B1C1 | 6 | 2 | 3 | 0 | 11 | 12 |
| L | 85 | M | A1B1C0 | 6 | 2 | 0 | 0 | 1 | 6 |

All patients were pathologically confirmed PD/PDD cases with a post-mortem delay between 4-7 hrs. y= year; F/M=female/male; ABC = ABC criteria according to Montine et al.[1]; Braak aSyn stage[2]; NFT= neurofibrillary tangle; Braak NFT stage[3]; Thal Phase[4]; CAA=Cerebral Amyloid Angiopathy[5]; ARTAG=Aging-related tau astrogliopathy[6]; LATE=Limbic-predominant age-related TDP-43 encephalopathy; PART=Primary Age-Related Tauopathy; AGD=Argyrophilic grain disease;[#]=LRP10 genetic variant; *The total numbers of inclusions per donor are not indicative of the total prevalence of each type of inclusion within each donor.

**Supplementary Table 2. Immunohistochemistry conditions for primary antibodies.**

| Antibody | Antibody target | Company information | Dilution from stock | Incubation conditions | Antigen retrieval conditions |
|---|---|---|---|---|---|
| LB509 | aSyn | Life Technologies (# 180215) | 1/500 | 1 hour, 37 °C | None |
| SYN-1 (Clone 42) | aSyn | BD Biosciences (# 15895639) | 1/100 | 4 hours, room temperature | 100% formic acid 10 mins, Tris-EDTA pH 9 at 100 °C for 30 mins |
| EP1536Y | $aSyn^{pS129}$ | Abcam (# 51253) | 1/1000 | overnight, 4 °C | None |
| 11a5 | $aSyn^{pS129}$ | Prothena (gift) | 1/100,000 | 1 hour, 37 °C | None |

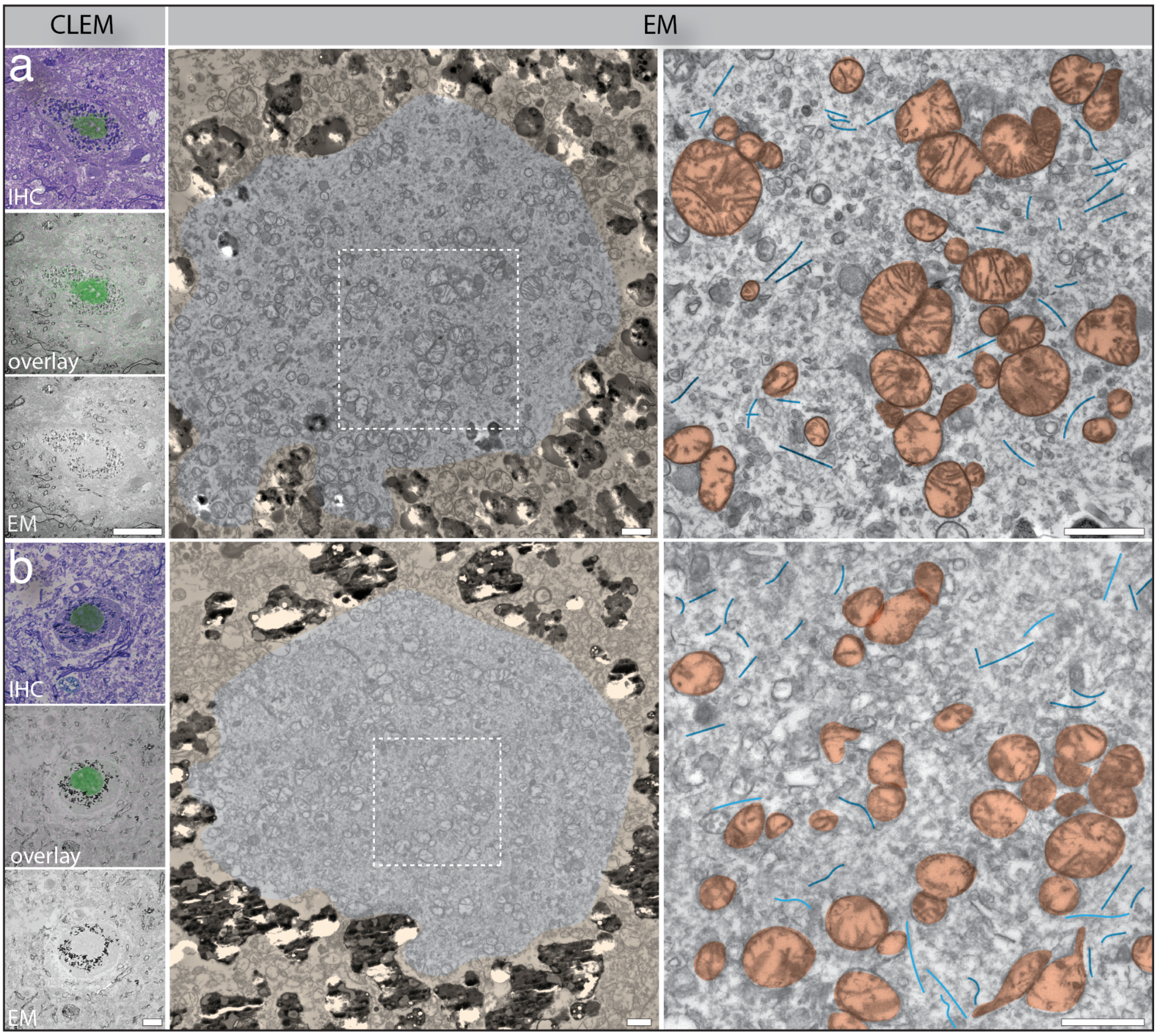


**Supplementary Figure 1. Further examples of pale bodies.** Large aggregates in neuromelanin containing neurons were localized by CLEM$^{IHC}$ and show aSyn immunostaining (a – SYN-1, BD Biosciences; b - 11a5, Prothena; green) overlaid on toluidine blue image from adjacent sections. Overlay panels show the correlated EM. The EM shows an ultrastructural arrangement of accumulated and clustered mitochondria (dark orange) interspersed amongst fibrillar material (blue). The tissue surrounding aSyn immuno-positive areas are false-coloured light orange for clarity. Scale bars: CLEM (a) 5 µm; CLEM (b) 10 µm; EM (center and right panels) 1 µm.

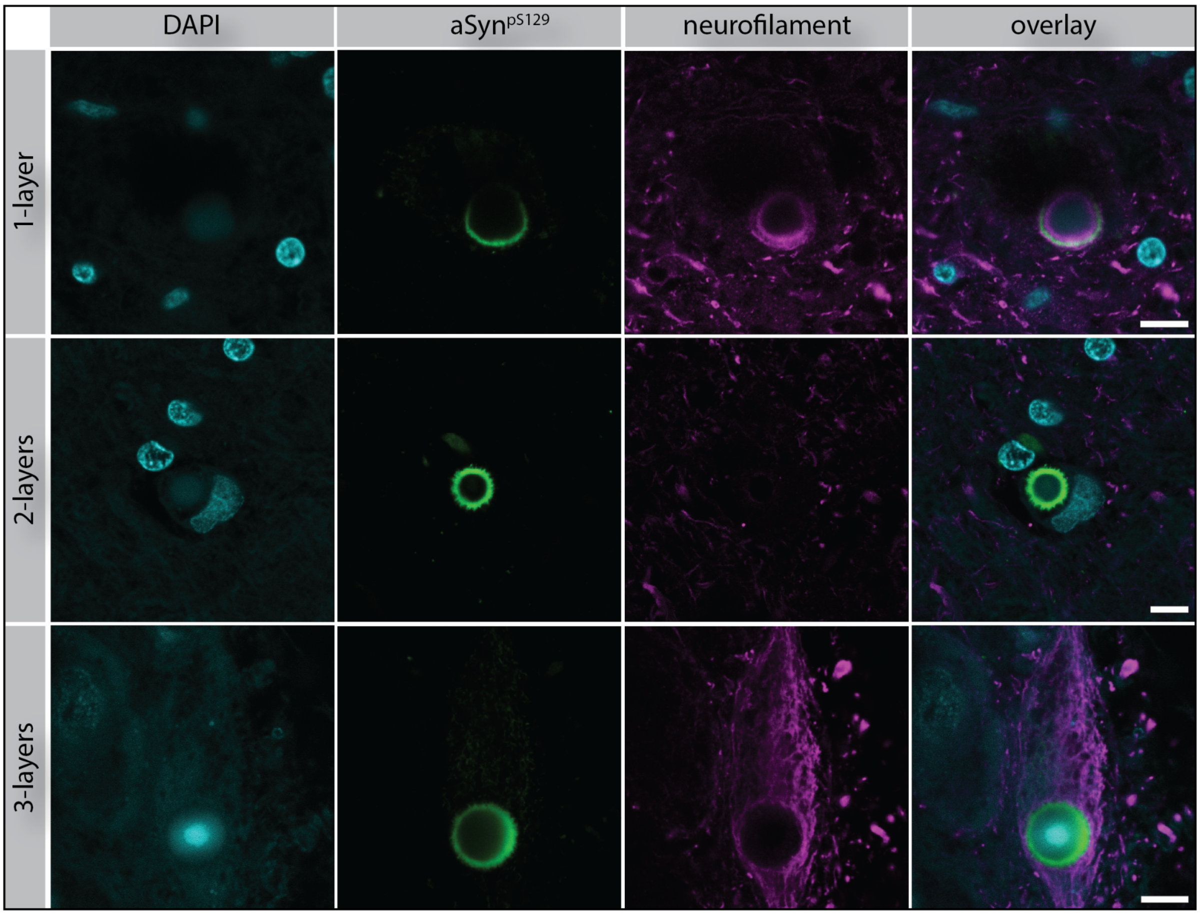


**Supplementary Figure 2: Neurofilament staining in Halo Lewy Bodies.** Raw fluorescent images from the central plane of the halo LBs shown in Figure 3. Immunostaining for cell nuclei (DAPI; cyan), aSyn$^{pS129}$(11a5, Prothena; green), and neurofilament (neurofilament H, Sigma; magenta) is shown. A double neurofilament ring surrounding the aSyn$^{pS129}$ halo is observed for the one-layer LB, a very faint neurofilament ring was observed for the two-layer LB, and the three-layer LB shows cellular neurofilament staining concentrated around the periphery of the LB. DAPI staining is present in the core of all inclusions. Scale bars: 10 µm.

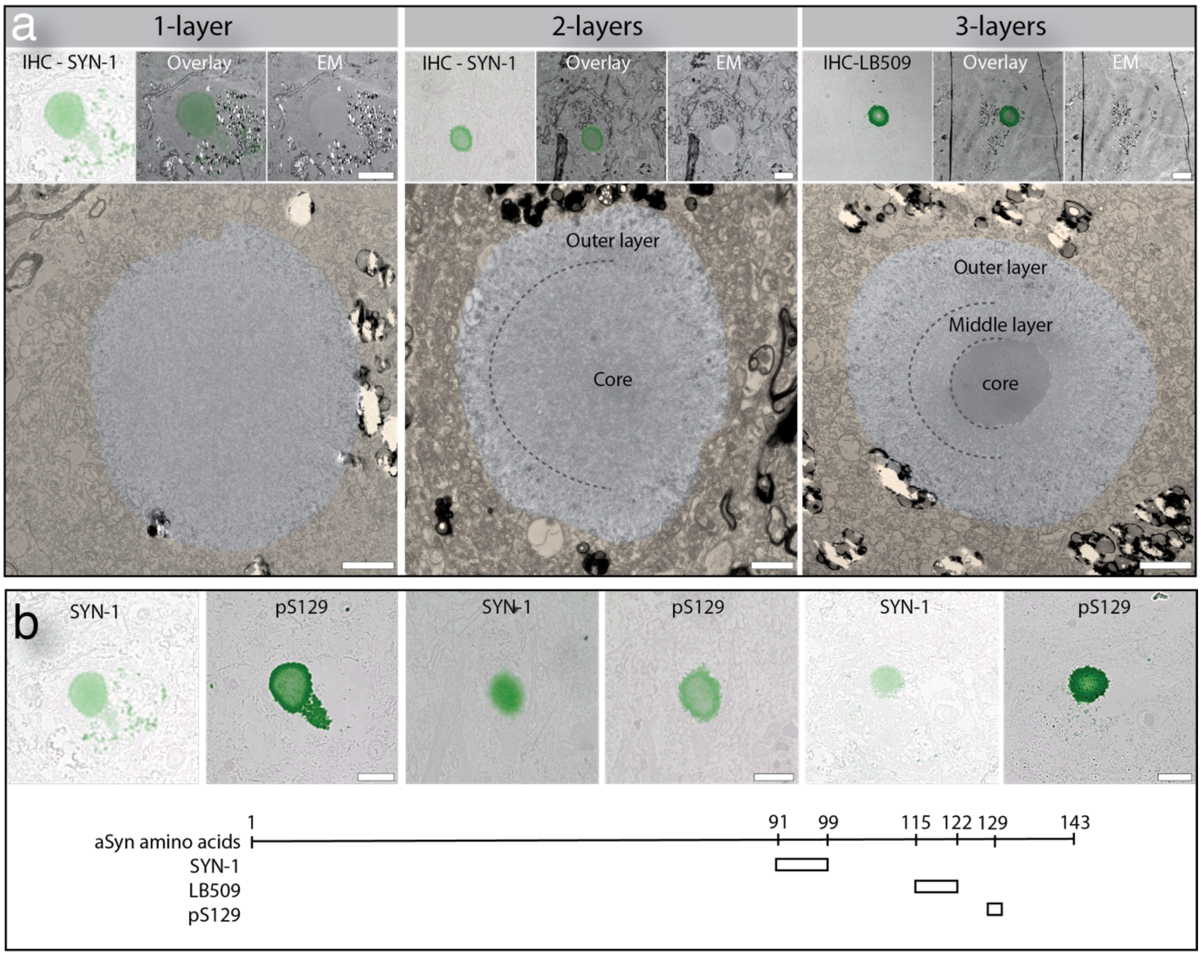


**Supplementary Figure 3: Halo Lewy Bodies localized within the same brain donor. (a)** Examples of one-, two- and three-layer halo LBs localized by CLEM$^{IHC}$ within the *substantia nigra* of donor B. The 1-layer LB is the same inclusion as shown in Figure 3a. aSyn immunostaining (one- and two- layer LBs: SYN-1, BD Biosciences; three-layer LB: LB509, Life Technologies; green) is shown. The tissue surrounding the LB is false-coloured orange for clarity. **(b)** Adjacent ultrathin sections for the LBs shown in (a) immunostained with SYN-1 (BD Biosciences) or aSyn$^{pS129}$ (EP1536Y, Abcam). The amino acid epitopes of aSyn recognized by each antibody used are shown. Scale bars: top panels in (a) 10 µm; bottom panels in (a) 2 µm; (b) 10 µm.

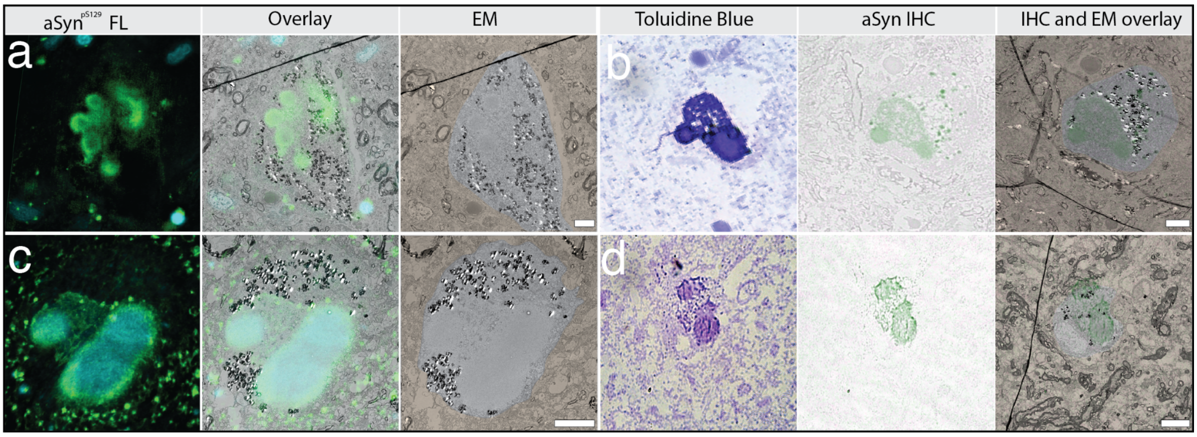


**Supplementary Figure 4: CLEM localization of multi-LB neurons.** aSyn immunostaining shown by fluorescence (a, aSyn$^{pS129}$ EP1536Y Abcam; c, 11a5, Prothena; green) or IHC (b, d – SYN-1, BD Biosciences; green) for inclusions shown in Figure 5. DAPI staining (cyan) is also shown for the CLEM$^{FL}$. Sections adjacent to the IHC were stained with toluidine blue for EM correlation. The inclusion in (c) was correlated using an adjacent grid to that shown in Figure 5c. The tissue surrounding the cell containing the inclusions is false-coloured light orange for clarity. Scale bars: 7 µm.

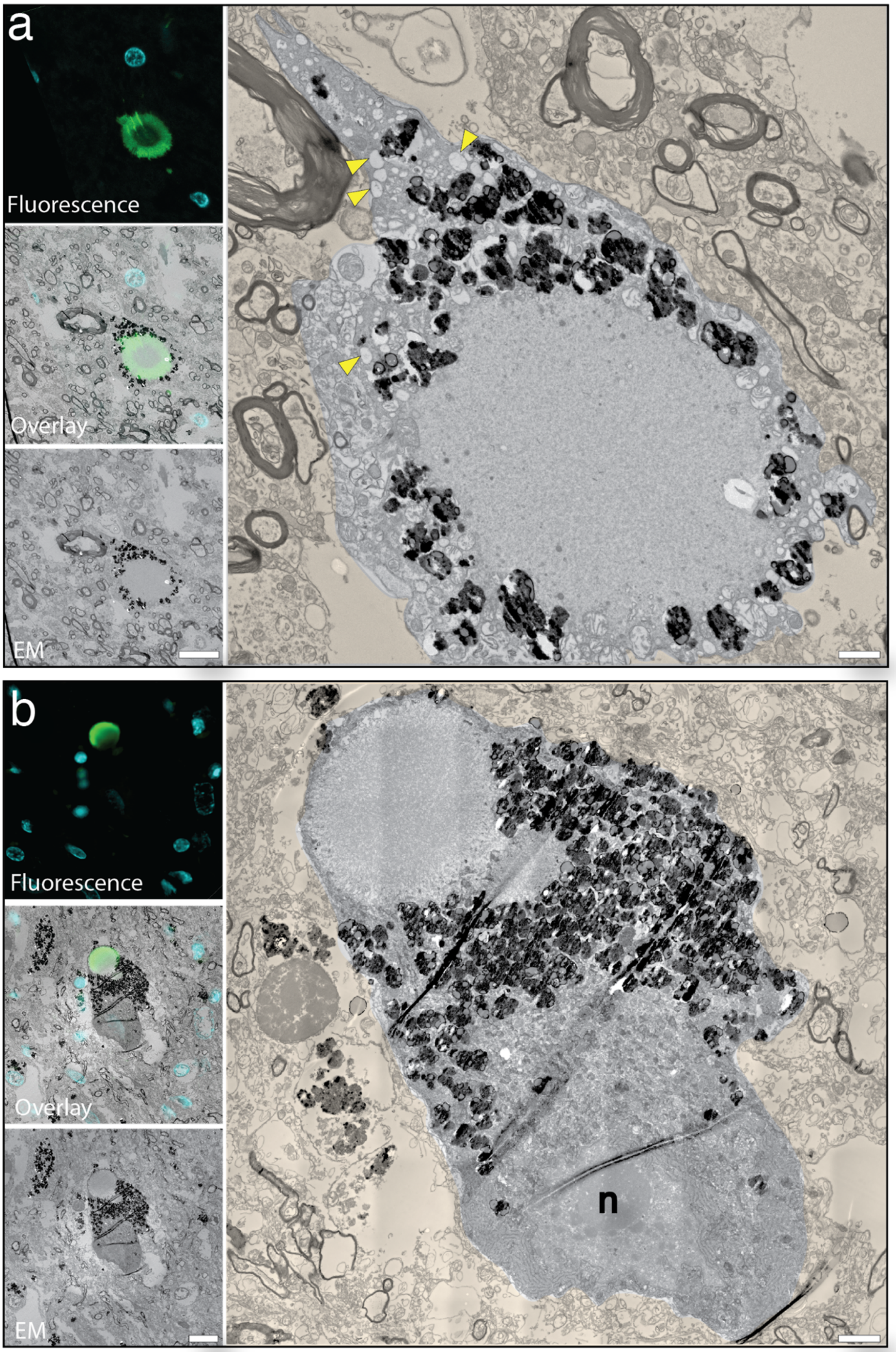


**Supplementary Figure 5: Degenerating or stressed neurons containing Lewy bodies.** The CLEM$^{FL}$ shows aSyn$^{pS129}$ (EP1536Y, Abcam; green) and DAPI (cyan). The neurons display morphological features consistent with degeneration or stress, including an electron dense nucleus (n) and cytoplasm, and frequent vacuoles (yellow arrowheads). The neuromelanin is visible in both neurons. The LB in (a) shows a 1-layer fibrillar ultrastructure. The LB in (b) shows a small 1-layer LB next to a larger 2-layer fibrillar LB, consisting of a peripheral region of loosely organized fibrils surrounding a denser fibrillar core. The tissue surrounding the cells are false-coloured light orange for clarity. Scale bars: CLEM 10 µm; TEM 2 µm.

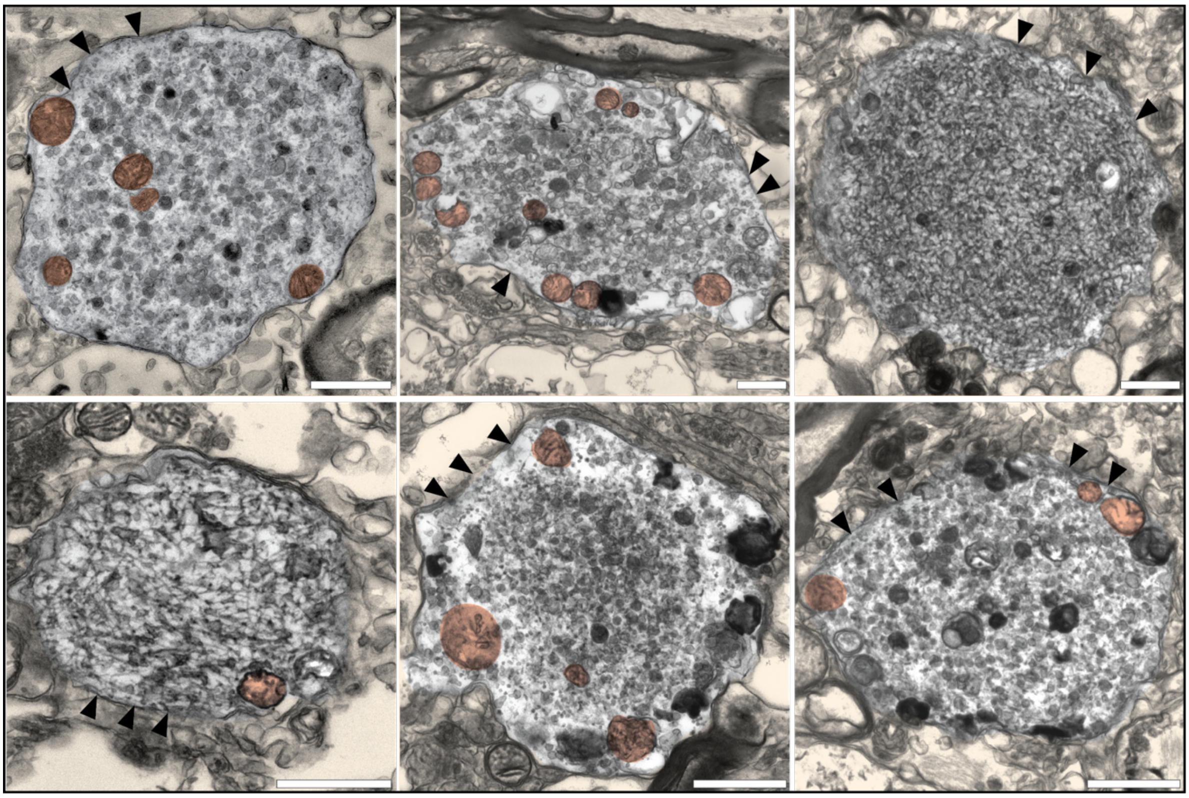

**Supplementary Figure 6: Small membranous only inclusions.** TEM images of compact neuritic aggregates < 5 µm showing densley accumulated membranous material with no evident fibrils. The surrounding neuritic membrane (black arrowheads) and internal mitchondria (dark orange). The aggregates were confirmed to be uniformly αSyn-immunopositive across their entire volume by CLEM$^{IHC}$ using LB509 (Life Technologies; data not shown). The tissue surrounding the cells are false-coloured light orange for clarity. Scale bars: 1 µm.

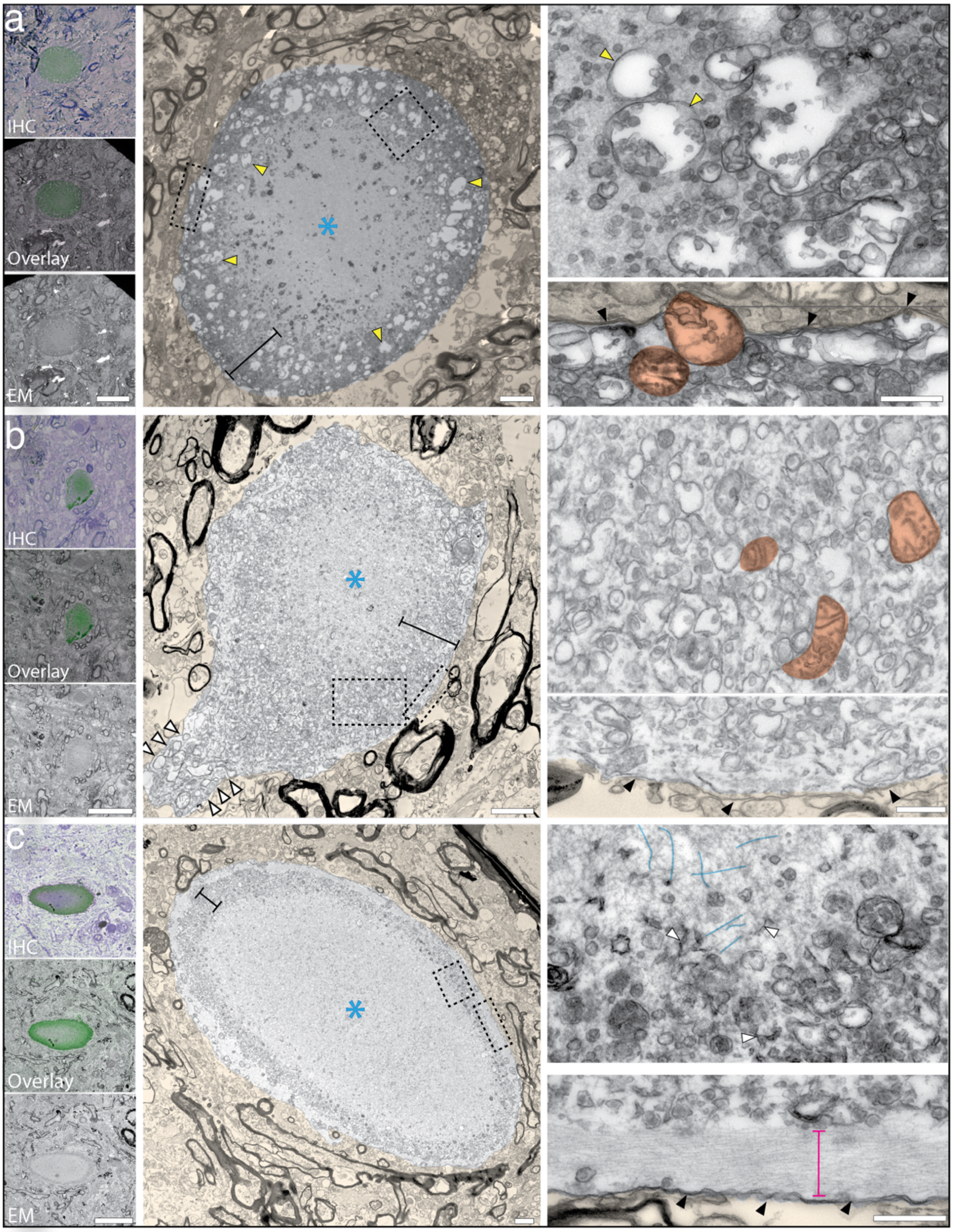


**Supplementary Figure 7: Further examples of the membranous halo fibrillar core ultrastructure.** The CLEM$^{IHC}$ shows aSyn immunostaining (a,b, LB509, Life Technologies; c, EP1536Y, Abcam; green) overlaid on toluidine blue images from adjacent sections. In the higher magnification EM images (black dashed boxes), the surrounding cytoskeleton filaments (magenta bracket) and neuritic membrane (black arrowheads) are indicated. Inclusions show varying membranous halo thickness (black bracket) surrounding a fibrillar core (blue asterisk), with a particularly thin and well-defined halo in (c). The membranous composition also varies between inclusions with (a) containing many vacuoles (yellow arrowheads). The neuritic membrane surrounding the inclusion in (b) can be seen to extend into a neuritic projection (magenta arrowheads). Mitochondria are coloured orange, examples of aSyn fibrils in the core of the inclusions are coloured blue. The tissue surrounding the inclusion has been false-coloured light orange for clarity. Scale bars: CLEM 20 µm; low mag TEM 2 µm; high mag TEM 500 nm.

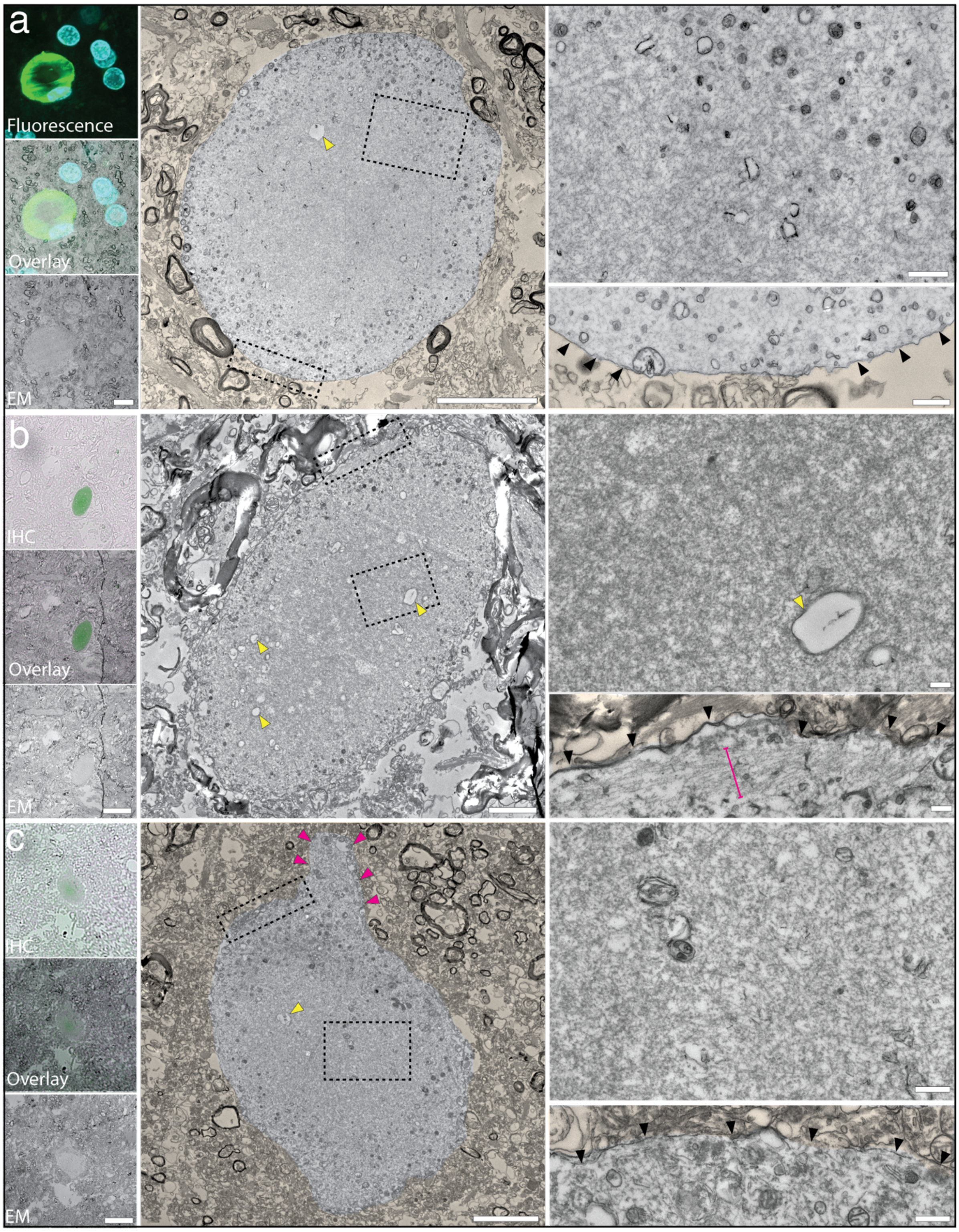


**Supplementary Figure 8: Predominantly fibrillar neuritic pathology.** The CLEM$^{FL}$ shows aSyn$^{pS129}$ (EP1536Y, Abcam; green) and DAPI (cyan); CLEM$^{IHC}$ shows aSyn (SYN-1, BD Biosciences; green). EM shows predominantly fibrillar inclusions with sparse peripheral membranous material. The neuritic membrane surrounding the inclusions are indicated with black arrowheads and can be seen to extend into a neuritic projection in (c) (magenta arrowheads). The inclusion shown in (b) has a ring of organized cytoskeleton between the neuritic membrane and the aSyn inclusion. Occasional vacuoles can be observed inside the inclusions (yellow arrowheads). The background tissue has been false-coloured light orange for clarity. Scale bars: CLEM - 20 µm; low mag EM- 5 µm, high mag EM -500 nm.

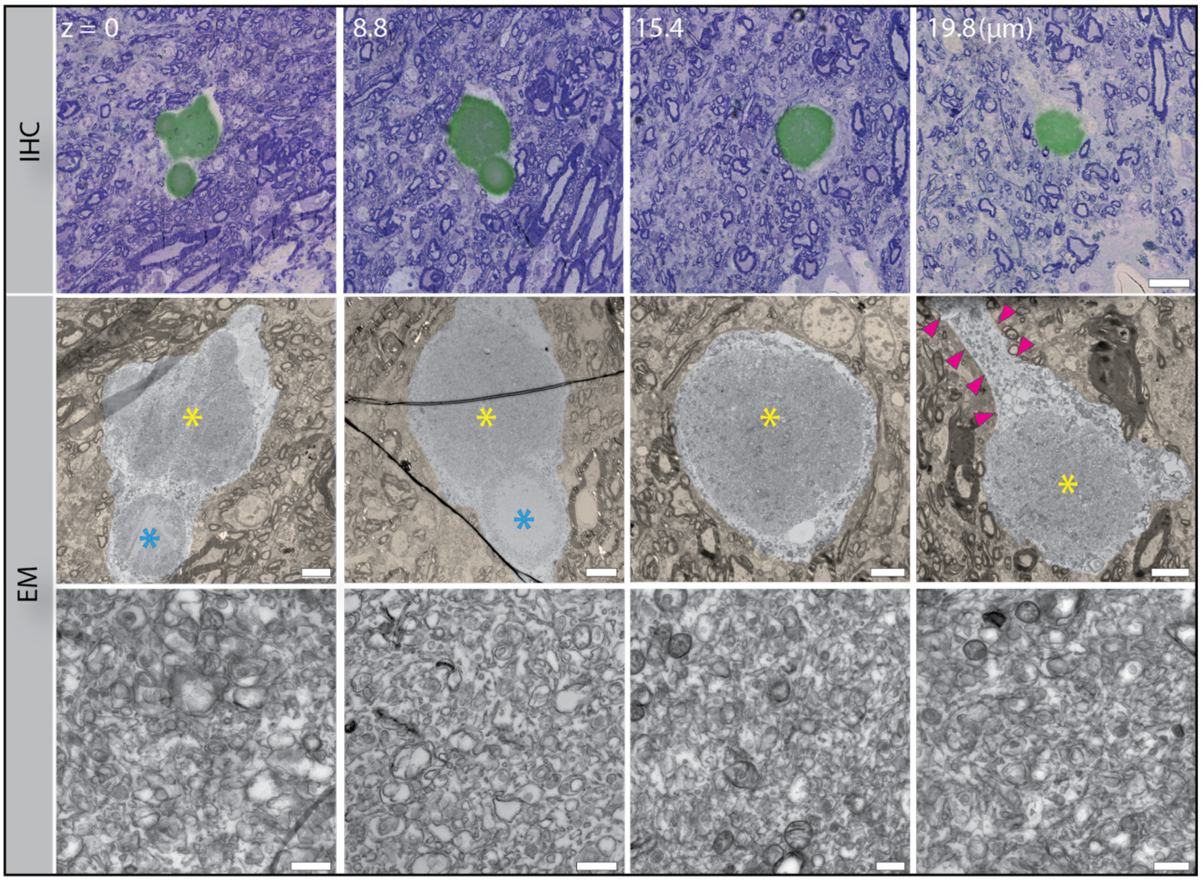


**Supplementary Figure 9**: **Multiple ultrastructures within the same neuritic inclusion.** The same inclusion from Figure 7 is shown across ~ 20 um of tissue along the z-axis. aSyn immunostaining (LB509; green) overlaid on toluidine blue images from the adjacent section is shown. High magnification EM throughout the inclusion shows densely accumulated membranous material, with no evidence of fibrils. At z =19.8 µm, the inclusion looks to be in a neuritic projection (magenta arrowheads). The inclusion at z =19.8 µm looks to be in a neuritic projection. Surrounding tissue is false-coloured light orange for clarity. Scale bars: low magnification EM 5 µm, high magnification EM 500 nm; IHC 20 µm.

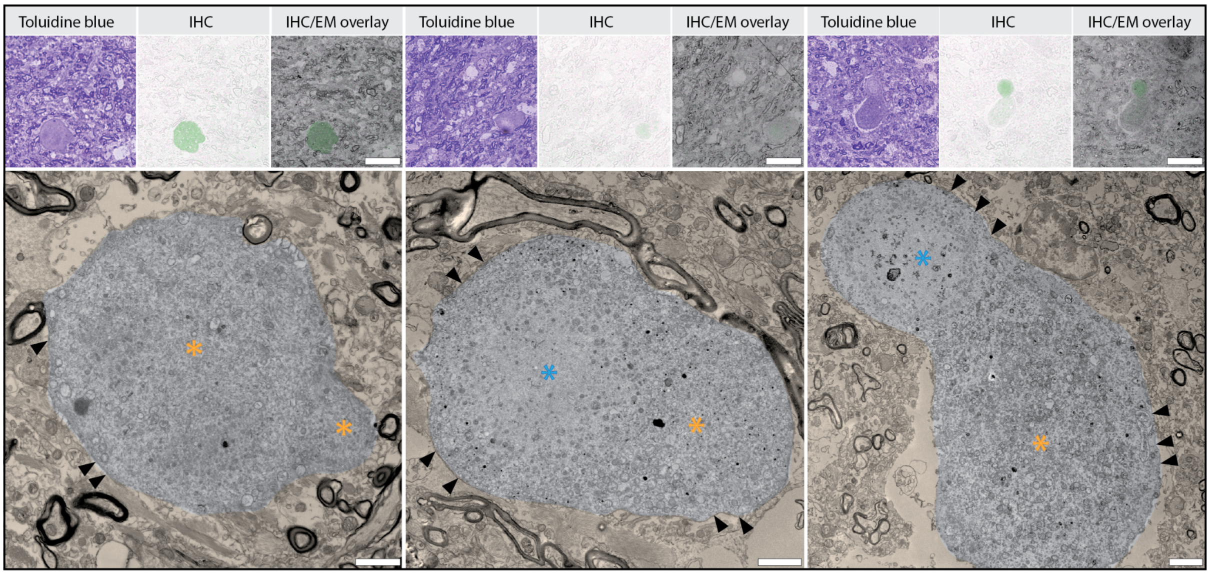


**Supplementary Figure 10: Further examples of merging/separating neuritic aSyn inclusions.** Multiple ultrastructures were observed within the inclusions including intermixed fibrils and membranes (orange asterisk) or a membranous halo, fibrillar core (blue asterisk). The CLEM$^{IHC}$ shows aSyn immunostaining (green; SYN-1, BD Biosciences). The toluidine blue image from the adjacent section is also shown. The neuritic membranes are indicated with black arrowheads. The tissue surrounding the aSyn immuno-positive region is false-coloured light orange for clarity. Scale bars: CLEM 20 µm; 2 µm.
